# G protein-coupled receptor signaling regulates ER-mitochondria contacts

**DOI:** 10.1101/2020.05.11.088815

**Authors:** Youngshin Lim, Il-Taeg Cho, Helmut G. Rennke, Ginam Cho

**Author notes:** Equal contribution (Youngshin Lim, Il-Taeg Cho).

## Abstract

Interactions between the endoplasmic reticulum (ER) and mitochondria (Mito) are crucial for many cellular functions, and their interaction levels change dynamically depending on the cellular environment. Little is known about how the interactions between these organelles are regulated within the cell. Here we screened a compound library to identify chemical modulators for ER-Mito contacts. Multiple agonists of G-protein coupled receptors (GPCRs), beta-adrenergic receptors (β-ARs) in particular, scored in this screen. Analyses in multiple orthogonal assays validated that GPCR activation promotes physical and functional interactions between the two organelles. Furthermore, we have elucidated potential downstream effectors mediating GPCR-induced ER-Mito contacts. Given the role of GPCRs in sensing various extracellular signals and subsequently eliciting appropriate cellular responses, our data provide significant insights into the roles of ER-Mito contacts in maintaining cellular homeostasis and in responding to physiological demands or stresses.

## Introduction

The membrane contacts between the ER and Mito are amongst the most extensively studied inter-organelle contacts (1). In yeast, ER-Mito contact sites are primarily tethered by the ER-Mito encounter structure (ERMES) complex (2, 3). In mammalian cells, the molecular identities of such complexes are less clear and the precise role of candidates is still debated (3). The functions of the ER-Mito contact sites include Ca^2+^ and lipid transfer, mitochondrial fission/fusion control, autophagosome/mitophagosome biogenesis and apoptosis control (1). The failure to maintain normal ER-Mito contacts is a common feature of neurodegenerative, cardiovascular, metabolic diseases, and cancer (4–7). For example, the disease-associated mutations in receptor expression enhancing protein1 (REEP1), linked to hereditary spastic paraplegia, result in decreased ER-Mito contact (8).

Accumulating evidence suggests that both abnormal increases and decreases in these contacts are detrimental to normal cellular physiology (4). If they are abnormally enhanced as in some Alzheimer’s disease models (4), the resulting Mito Ca^2+^ overload can lead to the opening of Mito permeability transition pores (mPTP) and subsequently to cell death (9). On the other hand, reduced or disrupted contacts, as in some models of amyotrophic lateral sclerosis (ALS) (4), are predicted to lower Mito ATP generation (9), which in turn is postulated to drive neuronal degeneration (10). Despite the increasing evidence implicating ER-Mito contacts in the pathogenesis of many diseases and highlighting their importance in maintaining normal cellular physiology, little is known about how the contacts are regulated.

GPCRs are well known to sense various extracellular stimuli, ranging from photons and ions to growth factors and hormones (11, 12). They transduce signals through heterotrimeric G proteins that consist of α, β, and γ subunits. Upon activation, the α subunit (G_α_), dissociated from the β/γ (G_βγ_) subunits, elicits subtype (G_s_, G_i_, G_q,_ and G_12_) specific downstream responses (13). Thus, the G-protein coupling preference of GPCRs determine the downstream pathways they activate. For instance, β-ARs, mainly coupled to G_s_, activates the pathway that relays a signal to adenylyl cyclase (AC) to generate cAMP whose downstream effectors include PKA (protein kinase A) and EPAC (exchange protein directly activated by cAMP) (13). Among the myriad of downstream events triggered by β-AR signaling, α-synuclein expression enhancement has been recently linked to an underlying drug mechanism of β-AR agonists, drugs shown to be associated with decreased risk in Parkinson’s disease (PD) and beneficial in several PD models (14, 15). The role of β-ARs in ER-Mito contacts is unknown.

We previously developed a split-*Renilla* luciferase reconstitution assay (referred to as split-Rluc assay hereafter) to quantitatively measure the level of physical coupling between ER and Mito (8, 16). Using this assay, we screened a compound library for ER-Mito contact enhancers and identified several agonists of GPCRs, mostly β-ARs. We verify that GPCR activation enhances physical as well as functional couplings between the two organelles. Uncovering the potential downstream effectors, our results provide novel insights into how GPCRs can modulate the ER-Mito coupling to elicit appropriate cellular responses. Furthermore, our successful compound screen demonstrates that the split-Rluc assay is an optimal, valid assay for screening, as it readily adapts to a high throughput detection in live cells without artifacts associated with fixation or complications related to image analysis.

## Results

### GPCR agonists were identified in a compound library screen for ER-Mito contact enhancers

We used the NIH Clinical Collection (NCC) library to screen for ER-Mito contact enhancers (Fig. 1). The HEK293T cells expressing split-Rluc constructs were treated with DMSO (control) or each drug (1 μM) (Fig. 1A, B), and the reconstituted split-Rluc activities (referred to as split-Rluc activities hereafter) in live cells were measured and normalized by the quantile method. From the initial screen, we identified 32 compounds (4.4% of the library) using the cut-off value of 3 standard deviation (SD) above the average of the control (Fig. 1C; Table S1). These compounds target GPCRs (18 drugs, 56% of the hits), glucocorticoid receptors (5 drugs, 16%; labeled as steroid), DNA topoisomerases/polymerases (5 drugs, 16%; labeled as DNA related), and others (3 drugs, 9%) (Fig. 1C, D, E; Table S1).

**Figure 1.**
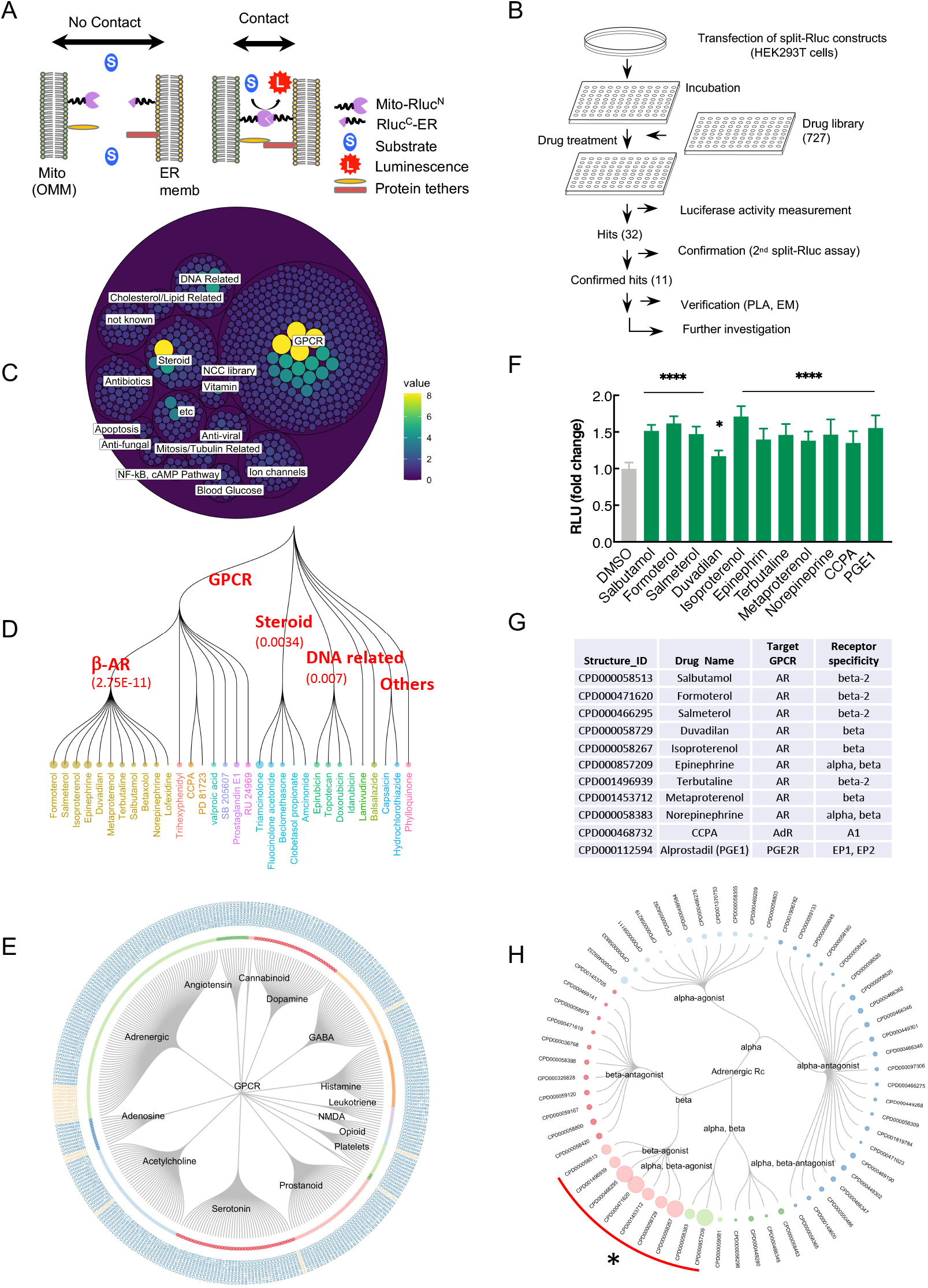
Compound library screening with split-Rluc assay identifies GPCR agonists. ***A***, *S*chematic of split-Rluc assay. ***B***, *S*chematic of screen workflow. ***C***, Initial screen results depicted with circular packaging dendrogram (each circle: each drug; circle size: normalized drug activity). Green to yellow circles indicate 32 hits. ***D***, Dendrogram of 32 initial hits. Circle size: normalized drug activity. Numbers in the parenthesis: false discovery rate (adjusted *p* values). ***E***, Circular dendrogram of all GPCR-associated drugs grouped into each pathway (Drug IDs with orange color: 18 hits). ***F***, Repeated split-Rluc assay in HEK293T cells treated with 11 hits targeting GPCRs. RLU (relative light unit): normalized luciferase activity (to the mean of DMSO). *n* = 3 experiments; 1-way ANOVA with Dunnett’s multiple comparisons test; ^*^*p* = 0.0135; ^****^*p* < 0.0001. ***G***, List of 11 confirmed hits targeting GPCRs. AR: adrenergic receptor; AdR, adenosine receptor; CCPA, 2-chloro-N^6^cyclopentyladenosine; PGE1, prostaglandin E1; PGE2R, PGE2 receptor. ***H***, Circular dendrogram of all AR-associated drugs grouped based on target specificity. Circle size: normalized drug activity; circle color: target specificity; red curve with asterisk (*): 9 hits targeting ARs.

Given the high percentage (56%) of the GPCR-linked drugs among the compounds that scored in our screen, we selected the 18 GPCR-linked hits to test for the replication of the results. Repeated split-Rluc assay confirmed that 11 compounds showed consistent and significant increase in split-Rluc activity (Fig 1F). Among these 11 compounds, nine were agonists of β-ARs while the other two, alprostadil (a.k.a. prostaglandin E1, PGE1) and 2-chloro-N^6^cyclopentyladenosine (CCPA), target prostaglandin and adenosine receptors, respectively (Fig. 1G). Considering only 3% (10 of 331) of the GPCR-linked drugs and 1.3% (10 of 727) of the total drugs in this library are β-AR agonists (Fig. 1E, H; Table S2, S3), these data indicate an enrichment of β-AR agonists in our screen. Interestingly, while 9 compounds are agonists of β-ARs (epinephrine and norepinephrine activate both α- and β-ARs), all the other AR-associated compounds such as agonists and antagonists of α-ARs or antagonists of β-ARs, did not score as contact enhancers (Fig. 1H; Table S3), suggesting a specific role of β-AR activation in contact promotion.

To verify the results of our screen, we focused on a β-AR agonist isoproterenol for further analyses. First, a dose response of the drug effect was examined by treating cells with different concentrations (10nM to 5 μM) of isoproterenol. A concentration-dependent increase in split-Rluc activity was detected in cells treated with isoproterenol, revealing a dose response of the drug effect (Fig. S1A). Next, we tested the ability of a β-AR antagonist to counteract the effect of the agonist. A β-AR antagonist sotalol abolished the effect of isoproterenol (1.00 ± 0.124, DMSO; 1.32 ± 0.102, isoprot; 1.08 ± 0.114, isoprot/sotalol) (Fig. S1B), confirming the drug activity is specifically mediated through β-AR pathway. Furthermore, we tested cell line specificity of the drug effect using HeLa, C2C12, and Neuro2a cells. HeLa and C2C12 cells were responsive to isoproterenol like HEK293T cells, while Neuro2a cells were not (Fig S1C). This is likely due to the differences in receptor availability in these cell lines; Neuro2a cells have little or no β-AR expression, while the other three cell lines express β-ARs albeit weaker level in HeLa cells (see Table S4 for mRNA-seq data in these cell lines).

To address whether the agonist-induced split-Rluc activity results from enhanced transcription of the reporter constructs, we measured the effect of isoproterenol in the presence/absence of the transcription inhibitor actinomycin D (Act D) (Fig. S1D). In the cells treated with Act D alone, split-Rluc activity was lowered. However, isoproterenol treatment counteracted the inhibitory effect of Act D (Fig. S1D), supporting that the isoproterenol-induced split-Rluc activity results from the elevated physical coupling between the two organelles and not just from increased transcription alone. In fact, the expression level of the two split fragments (Rluc^C^-ER and Mito-Rluc^N^) were comparable between the vehicle- and isoproterenol-treated cells (Fig. S1E). Our data also indicate that the half-life of each fragment is relatively short (approximately 3 hrs for Mito-Rluc^N^; 5 hrs for Rluc^C^-ER) (Fig. S1F).

### GPCR agonists increase physical contacts between ER and mitochondria

As described above, we identified three classes of compounds in our screen: agonists of the β-AR, adenosine receptor, and prostaglandin receptor. We selected one member from each class to further investigate their potential roles in ER-Mito contacts: isoproterenol (PubChem CID 3779), a β-AR agonist; CCPA (PubChem CID 123807), adenosine A1 receptor agonist; and PGE1 (PubChem CID 5280723), which targets prostaglandin E2 receptor subtypes EP1 and EP2.

To corroborate our split-Rluc assay results and validate the drug effect on ER-Mito contacts, we employed three independent assays. First, a proximity ligase assay (PLA), which detects the proximity of the two proteins in the cell with specific primary antibodies and oligonucleotide-linked secondary antibodies (17). When SEC61 (ER protein) and TOM20 (mitochondrial protein) are in sufficient proximity, the interaction between these two proteins—i.e., membrane-membrane apposition between ER and Mito—generates a fluorescent product (red) that can be detected *in situ* (Fig. 2A). Isoproterenol-, CPA-, or PGE1-treated cells showed significant increase in PLA fluorescent signal, demonstrating that these drugs indeed facilitate ER-Mito contacts (Fig. 2A, B and Fig. S2A). Of note, different treatment time of isoproterenol resulted in different level of responses in PLA signal (Fig. S2B), suggesting a dynamic nature of the drug effect on ER-Mito contacts.

**Figure 2.**
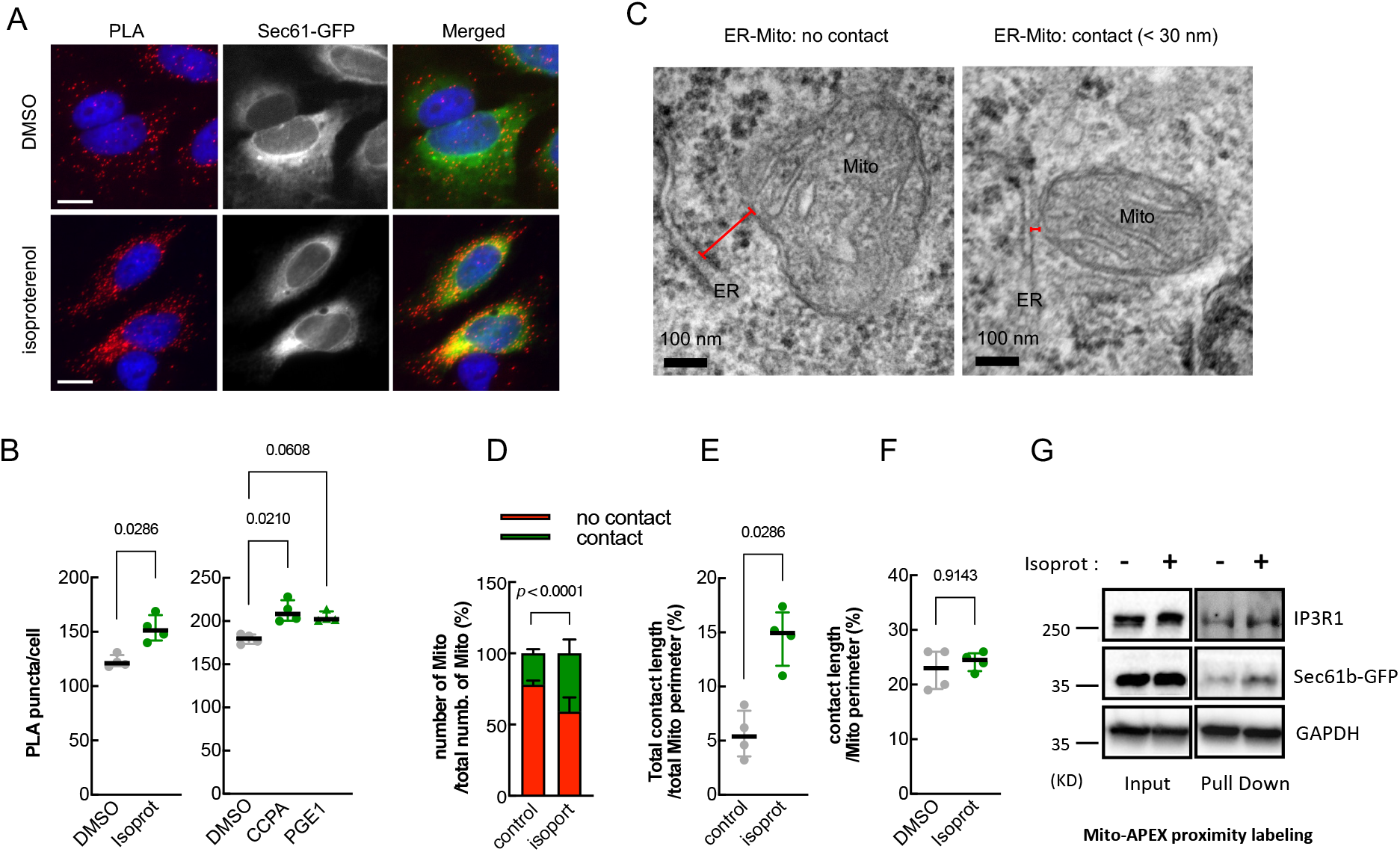
GPCR agonists enhance physical contacts between ER and Mito. ***A***, Representative images of PLA in HeLa cells treated with DMSO or isoproterenol (1 μM). PLA signal (red) indicates a close contact between ER and Mito. Sec61-GFP (green): to label ER; DAPI (blue): nucleus. ***B***, Quantifications of PLA signal as shown in A (left) and S2A (right) (*n* = 4 experiments with 77-97 cells each; Mann-Whitney test (left); Kruskal-Wallis test with Dunn’s multiple comparisons test (right). ***C***, Representative TEM images of Mito (HEK293T cells) showing no contact (left) or a close contact (right) with ER membrane. Red lines: the distance between ER and Mito membrane. ***D***, Quantitative analysis of the percentage of the number of Mito showing contact or no contact with ER (*n* = 3 experiments with 346, 405 Mito for control, isoprot; Chi-square test). ***E-F***, The percent of total contact length divided by the total Mito perimeter measured among all mitochondria (E) (*n* = 4 experiments with 122, 171 mitochondria for control, isoprot; Mann Whitney test), and the percent of contact length divided by Mito perimeter using only Mito showing a close contact with ER (F) (*n* = 4 experiments with 27, 95 mitochondria for control, isoprot; Mann Whitney test) in TEM images. ***G***, Representative images of Western blots of biotinylated proteins proximity-labeled by Mito-APEX (for 3min) followed by streptavidin affinity pull down, in control- vs isoprot-treated HEK293T cells. GAPDH used for control. Data in B E, F: median with interquartile range; Data in D: mean ± SD. Numbers at the top of each graph (here and in all other figures): *p* values (if ≤ 0.05, significant). Scale bars: 10 μm (A), 100 nm (C).

Next, we assessed the contacts at the ultrastructural level by transmission electron microscopy (TEM). The cells treated with or without isoproterenol were fixed and analyzed by TEM. The relative number of mitochondria closely juxtaposed (< 30 nm) to ER over the total number of mitochondria in isoproterenol-treated cells was significantly higher (DMSO, 22.5%; isoproterenol, 41.7%; *p* < 0.0001; Chi-square test) (Fig. 2C, D). Furthermore, the isoproterenol-treated cells showed a significant increase in total contact coverage, calculated as the percent of the total contact length divided by the total Mito perimeter measured among all mitochondria (5.6 ± 2.1%, control; 14.6 ± 2.7%, isoprot; *p* = 0.0286) (Fig. 2E). Interestingly, when only the mitochondria having a tight coupling with ER were compared, the percent of contact length did not show a statistically significant difference (Fig. 2F). We speculate that isoproterenol might increase the proximity of the ER and Mito (contact width) or the number of contacts, with little or no effect on contact length.

Finally, the ascorbate peroxidase (APEX)-proximity labeling method was employed which biotinylates proteins within 20 nm proximity of the enzyme. Mito-APEX, targeted to Mito outer membrane, can label and identify ER proteins in close proximity to the Mito (16). The cells transfected with mito-APEX were treated with DMSO or isoproterenol, and the biotinylated proteins (by incubating with biotin-phenol and with H_2_O_2_) were affinity-purified using streptavidin-magnetic beads and subjected to Western blot analysis. In these affinity purified samples (which were within 20nm proximity to the Mito membrane) prepared from the isoproterenol-treated cells, we found that ER proteins IP3R1 and SEC61b were increased, but not cytosolic protein GAPDH (Fig. 2G). Together, these data from three independent assays corroborate our results with the split-Rluc assay, strongly supporting that GPCR agonists enhance physical contacts between the ER and Mito.

### *GPCR agonists facilitate* mitochondrial *Ca*^*2+*^ *uptake and bioenergetics*

We next investigated if drug-induced physical contacts created functional coupling between the ER and Mito. Given the role of ER-Mito contact sites as hotspots for Ca^2+^ influx to Mito (9), we explored if agonist treatment increases Mito Ca^2+^ influx, using a fluorescent Ca^2+^ sensor mito-R-GECO1 (18). The cells were pre-treated with DMSO, isoproterenol, CCPA, or PGE1, and the Mito Ca^2+^ uptake was measured every 2.5 seconds in response to histamine-stimulated ER Ca^2+^ release (Fig. 3A). Upon ER Ca^2+^ release, all three drugs elevated Mito Ca^2+^ uptake (Fig. 3A), resulting in a higher maximum peak (Fig. 3B) and faster uptake rate (Fig. 3C). Also, when ionomycin (Ca^2+^ ionophore) was used to trigger cytoplasmic Ca^2+^ influx, a similar increase in Mito Ca^2+^ uptake was observed (Fig. 3D). Together these data support that these agonists enhance, not only physical, but also functional coupling between the ER and Mito.

**Figure 3.**
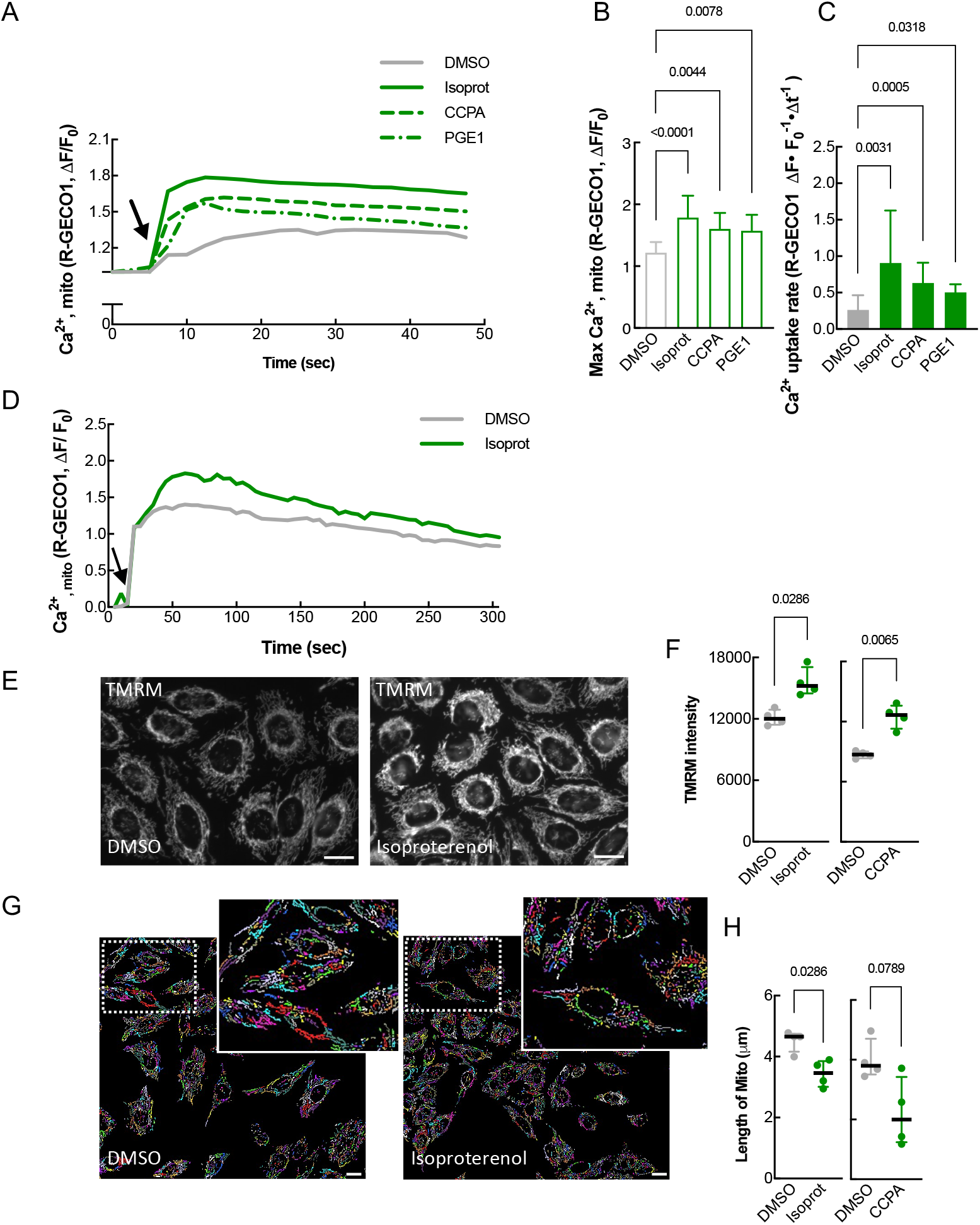
GPCR agonists enhance functional coupling between ER and Mito. ***A***, Mito Ca^2+^ uptake in response to histamine (100 μM). Mito-R-GECO1 fluorescent intensity change averaged over multiple cells (is plotted every 2.5 sec in DMSO, isoprot, CCPA, and PGE1 treated cells. Arrow: histamine addition. ***B***, Maximum peak of Mito Ca^2+^ uptake (as in A). One-way ANOVA with Dunn’s multiple comparisons test. ***C***, Mito Ca^2+^ uptake rate (as in A; 7.5 to 12.5 sec). Kruskal-Wallis test with Dunn’s multiple comparisons test. ***D***, Mito Ca^2+^ uptake (HeLa cells) in response to ionomycin (3 μM). Mito-R-GECO1 fluorescent intensity change is plotted every 5 sec. ***E***, Representative images of TMRM signal (for Mito membrane potential) (HeLa cells). ***F***, Quantification of TMRM intensity as in E (left) or S2B (right). (Left) *n* = 4 with 52 cells each; unpaired two-tailed *t*-test with Welch’s correction. (Right) *n* = 4 with 72-100 cells each; Kruskal Wallis test with Dunn’s multiple comparison test. ***G***, Representative images of Mito (converted in Image Analyst MKII software) used for length comparison (HeLa cells, 1 μM drug). ***H***, Quantification of Mito length as in G (left) or S2C (right). (Left) *n = 4* with 616 (DMSO) or 948 (Isoprot) Mito for each experiment; Mann Whitney test. (Right) *n = 4* with 278 (DMSO) or 445 (CCPA) Mito for each experiment; Kruskal Wallis test with Dunn’s multiple comparisons test. Each drug: used at 1 μM final conc. Data in B, C: mean ± SD; data in F, H: median with interquartile range. Scale bars: 10 μm.

ER-Mito coupling contributes to Mito bioenergetics by supplying Ca^2+^ (19–21), required for the function of proteins in the tricarboxylic acid (TCA) cycle and oxidative phosphorylation (21–23). We thus investigated the effect of the agonists on Mito bioenergetics. The Mito membrane potential, an indicator of increased mitochondrial metabolism (24), was measured with the tetramethyl rhodamine methyl ester (TMRM) fluorescence dye (Fig. 3E, F). Isoproterenol- or CCPA-treatment significantly increased the intensity of the TMRM signal (Fig. 3E, F and Fig. S2C). These findings imply that drug-induced ER-Mito contacts enhance Mito metabolism probably by facilitating ER-to-Mito Ca^2+^ transfer. Interestingly, isoproterenol led to a reduction in average Mito length in cells (Fig. 3G, H and Fig. S2D). Given that ER-Mito contacts are known to mark Mito fission sites (25), our data raise the interesting possibility that ER-Mito contact enhancement by GPCR activation may facilitate Mito fission events.

### cAMP/EPAC mediates β-AR signal-induced ER-Mito contacts

To gain insight into the downstream factors of the GPCR signaling involved in ER-Mito contact modulation, we evaluated the involvement of the G_s_ pathway because G_s_ is the primary G_α_ coupled to β-AR (Fig. 4A) (13). First, we examined the G_s_ pathway activation using a cAMP response element (CRE) reporter gene assay (a proxy for cAMP measurement). Compared to DMSO, isoproterenol increased CRE reporter activity in cells (fold change: 4.79 ± 1.66) (Fig. S3A), confirming G_s_ pathway activation by isoproterenol. Interestingly, when β2-AR itself was overexpressed, it induced much higher CRE reporter activity (fold change: 249.60 ± 19.90) (Fig. S3B) than isoproterenol treatment, arguing that the isoproterenol-mediated β-AR stimulation in our assay is not as strong as a full activation via receptor overexpression.

**Figure 4.**
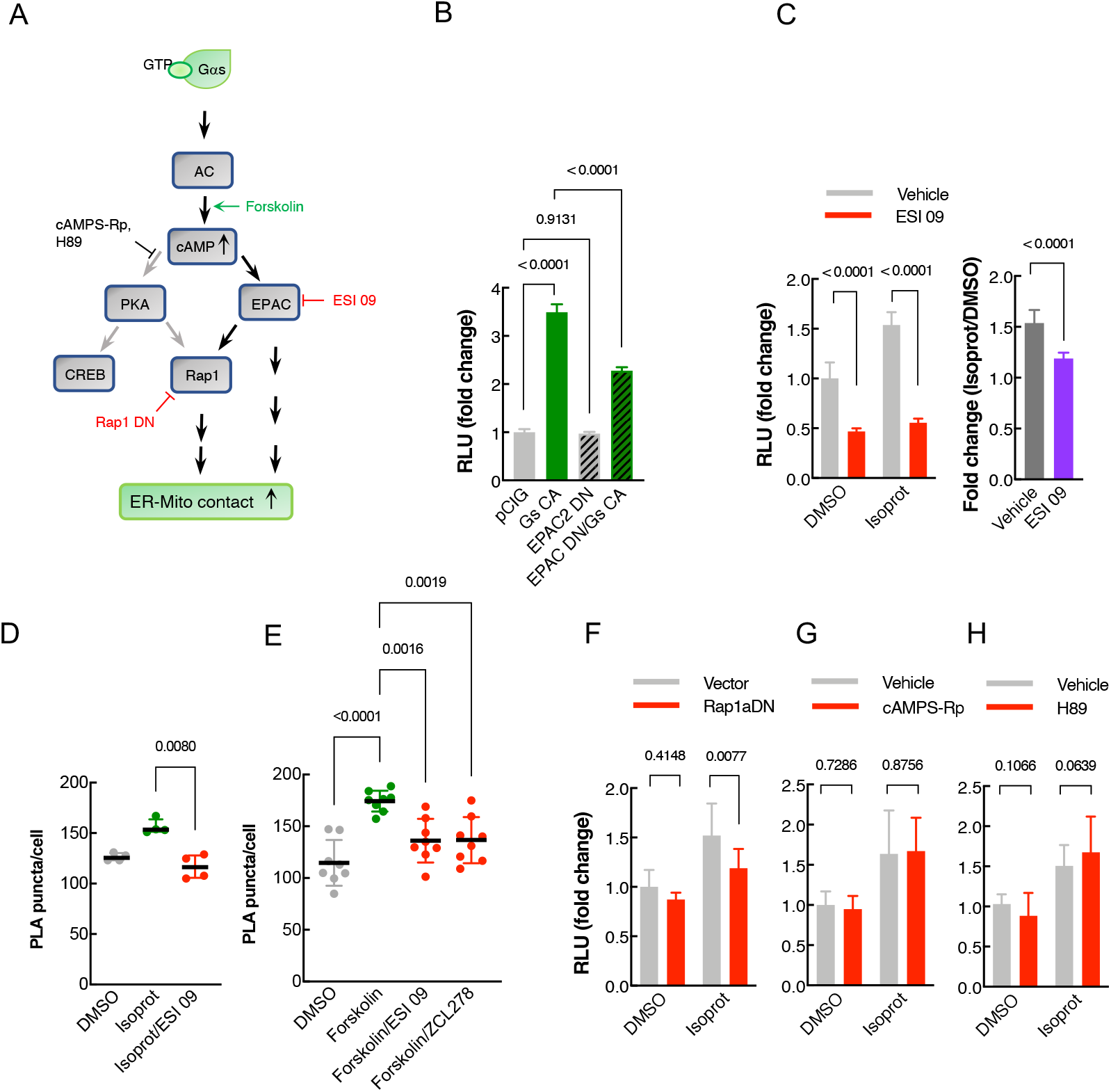
Ga/cAMP/EPAC pathway mediates β-AR signal-induced ER-Mito contacts. ***A***, Schematic of the simplified G_s_ and G_i_ pathway. Arrows: downstream pathways involved (black) or not-involved (grey) in β-AR-mediated contact modulation. Pathway inhibitors labeled with red (repressing) or black (not repressing) depending on their ability to repress agonist-induced contacts; activator (forskolin): green. ***B***, Split-Rluc assay in HEK293T transfected with pCIG (vector expressing GFP), G_s_ CA, EPAC DN, EPAC DN with G_s_ CA. *n* = 8; One-way ANOVA with Tukey’s multiple comparisons test. ***C***, (Left) Split-Rluc assay in H293T cells treated with vehicle vs ESI 09 (5 μM) in the presence of DMSO or isoprot. *n* = 8; 2-way ANOVA with Sidak’s multiple comparisons test. (Right) Fold change of isoproterenol-induced luciferase activity (isoprot/DMSO) in the presence of vehicle vs. ESI 09 (using the data sets from the left graph); two-tailed *t*-test. ***D-E***, Quantification of PLA in HeLa cells treated with DMOS, isoprot, or isoprot with ESI 09 (D) (*n* = 4 with 64 (DMSO), 107 (isoprot), 70 (Isoprot/ESI 09) cells each; Kruskal Wallis test with Dunn’s multiple comparisons test), or with DMSO, forskolin, forskolin with ESI 09, or forskolin with ZCL 278 (*n* = 8 with 117-167 cells each; one-way ANOVA test with Sidak’s multiple comparisons test). ***F-H***, *S*plit-Rluc assay in HEK293T cells transfected with vector or Rap1a DN (G, *n* = 8), or treated with vehicle or PKA inhibitors cAMPS-Rp (10 μM; *n* = 20) (H) and H89 (10 μM; *n* = 15) (I), in the presence of DMSO or isoprot; 2-way ANOVA with Sidak’s multiple comparisons test. All data except D and F: mean ± SD; data in D and F: median with interquartile range.

Next, we asked if G_s_ pathway activation influences the level of ER-Mito contacts. A constitutively active construct for G_s_ (G_s_ CA), which activates the signaling cascade even in the absence of ligand (26), augmented split-Rluc activity (Fig. 4B). Corroborating these results, an adenylyl cyclase (AC) activator forskolin (thus increasing cAMP level) enhanced PLA signal (Fig. 4E). Together these results demonstrate that G_s_/AC/cAMP pathway activation leads to an increase in ER-Mito contacts.

Because cAMP can transduce signal through two different downstream effectors, EPAC and PKA, we next examined which of these factors were involved in mediating the signal for contact regulation. Using two different tools, a dominant negative (DN) construct and a pharmacological inhibitor (ESI 09), we determined if EPAC activity is required for G_s_/AC/cAMP pathway-induced contacts. The EPAC DN construct significantly reduced G_s_ CA-induced split-Rluc activity (fold change: 3.49 ± 0.164, G_s_ CA; 2.28 ± 0.074, G_s_ CA/EPAC DN; *p* < 0.0001) without affecting basal level of split-Rluc activity (fold change: 1.00 ± 0.069, pCIG (vector); 0.97 ± 0.035, EPAC DN; *p* = 0.9131) (Fig. 4B). Similarly, a chemical inhibitor of EPAC, ESI 09, caused a significant reduction in isoproterenol-induced split-Rluc activity (Fig. 4C, left). Because it also reduced the split-Rluc activity in DMSO-treated cells (Fig. 4C, left), we compared the fold changes of the isoproterenol-induced activity (over DMSO) between the vehicle- vs. ESI 09-treated cells and found a statistically significant difference (1.54 ± 0.128, vehicle; 1.19 ± 0.057, ESI 09; *p* < 0.0001) (Fig. 4C, right). Together with EPAC DN data, these results confirm the requirement of the EPAC for the isoproterenol-induced contacts, although ESI-09 appears to affect basal level contacts as well. Corroborating these results, the PLA also demonstrated a similar inhibitory effect of ESI 09 (Fig. 4D). Consistent with these data, ESI 09 also blocked forskolin-induced PLA signal enhancement (forskolin increases cAMP by stimulating adenylyl cyclase) (Fig. 4E). Of note, we found that RAP1A (EPAC downstream effector) is also involved, as RAP1A DN decreased the isoproterenol response (Fig. 4F), albeit its effect is not as robust as EPAC inhibition (Fig. 4D); this implicates that EPAC-mediated effect is only partially transmitted through RAP1A.

In contrast to EPAC, the inhibition of PKA activity by Rp-cAMPS and H-89 (Fig. 4G, H), or of MEK activity by PD 032509 (Fig. S3C) did not show any blocking effect on isoproterenol response. Together these results support that EPAC, but not PKA, is a downstream factor mediating β-AR signaling for ER-Mito contacts. In accordance with our data, a recent study has demonstrated that EPAC is necessary for increasing the interaction between VDAC1, IP3R1, and GRP75, a well-established complex at ER-Mito contact sites (27).

### Elevated cytosolic Ca^2+^ participates in β-AR signal-induced ER-Mito contacts

In addition to cAMP, cytosolic Ca^2+^ level is also known to increase upon β-AR activation by isoproterenol (Fig. 5A) (28). We thus set out to evaluate the potential role of cytosolic Ca^2+^ in isoproterenol-induced ER-Mito interactions. First, we ensured an increase in cytoplasmic Ca^2+^ level upon isoproterenol application to the cells by using fluorescent Ca^2+^ sensor GCaMP6s (Fig. 5B). Next, we investigated if intracellular Ca^2+^ increase itself could enhance ER-Mito interactions. For this, we assessed the effects of the three different drugs, histamine, ionomycin and cyclopiazonic acid (CPA), all of which are known to elevate cytosolic Ca^2+^ albeit by distinct mechanisms (28, 29). Histamine stimulates cytosolic Ca^2+^ release from the ER through IP3 receptor activation (28); ionomycin, a membrane permeable Ca^2+^ ionophore, allows both Ca^2+^ release from intracellular stores and extracellular Ca^2+^ influx to the cytoplasm (28); and CPA, a sarco-endoplasmic reticulum Ca^2+^-ATPase (SERCA) inhibitor, blocks the uptake of Ca^2+^ ions back to the sarcoplasmic reticulum (SR)/ER lumen after their release into the cytosol, thus accumulating Ca^2+^ in the cytoplasm (29). All three drugs elevated Split-Rluc activity (Fig. 5C), suggesting that increasing cytoplasmic Ca^2+^ concentration is sufficient to enhance ER-Mito contacts. We corroborated these findings with the PLA results; histamine or ionomycin treatment increased PLA signal (Fig. 5D).

**Figure 5.**
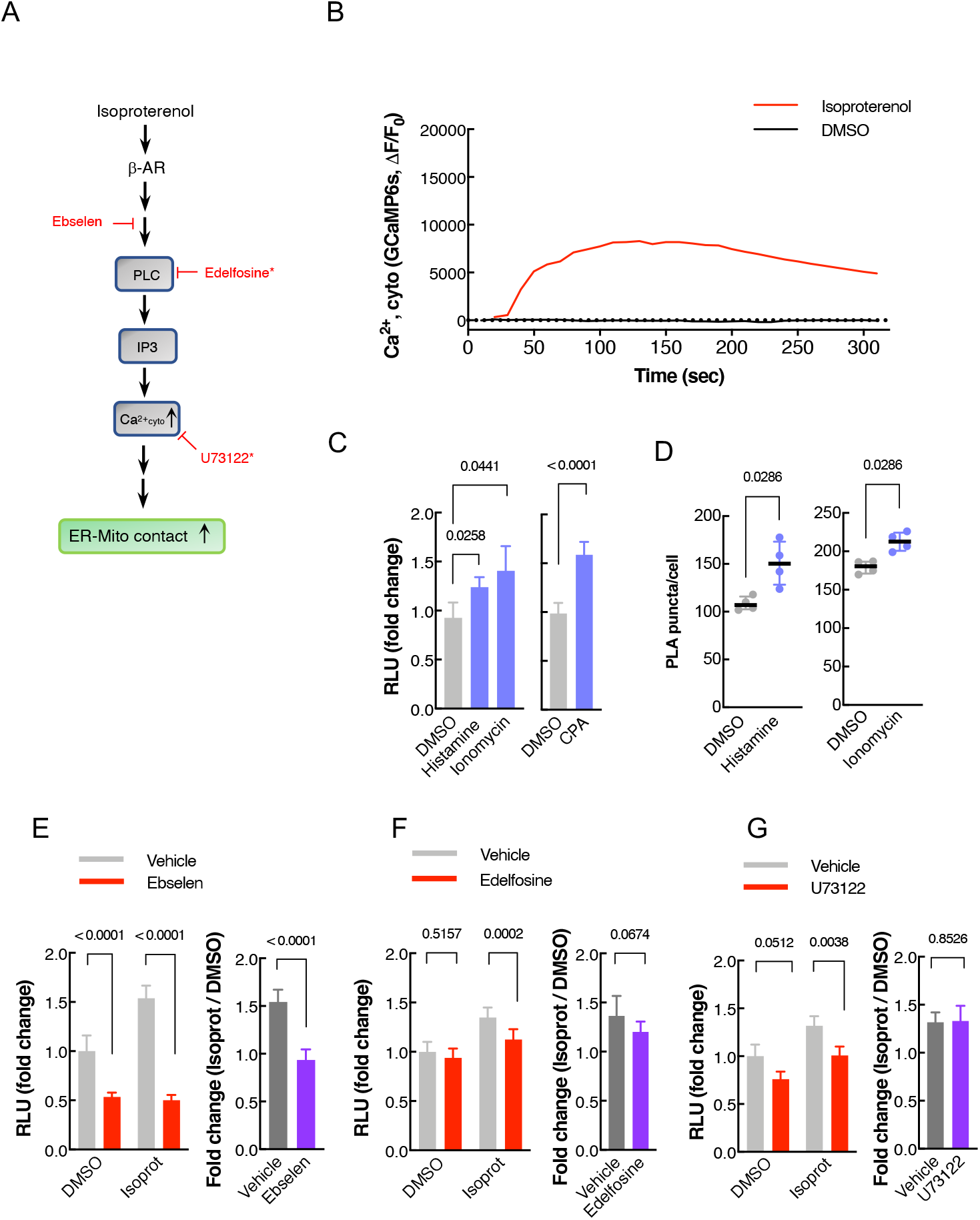
Cytoplasmic Ca^2+^ participates in β-AR signal-induced ER-Mito contact. ***A***, Schematic of the simplified β-AR pathway leading to cytosolic calcium increase. Chemical inhibitors: red. Asterisk (*): drugs with negligible effect on isoproterenol-induced ER-Mito coupling. IP3: inositol triphosphate; PLC: phospholipase C; GEFs: Guanine nucleotide exchange factors; ***B***, Cytoplasmic Ca^2+^ level measured by GCaMPs fluorescence intensity fold change (ι1F/F_0_) in HeLa cells treated with DMSO or isoprot (1 μM) every 2.5 sec (averaged over 16 cells). ***C***, Split-Rluc activities in DMSO, histamine (100 μM), ionomycin (1 μM), or CPA (10 μM) treated HEK293T cells. 1-way ANOVA with Dunn’s multiple comparisons test (left; n = 6); two-tailed *t*-test (right, *n* = 9). ***D***, Quantification of PLA in HeLa cells (*n* = 4 with 66, 57, 97, 87 cells for DMSO, histamine, DMSO, ionomycin; Mann Whitney test). ***E-G***, (Left) Split-Rluc assay in HEK293T cells treated with vehicle or indicated drugs (ebselen, 50 μM; edelfosine, 40 μM; or U73122, 5 μM), in the presence of DMSO or isoproterenol (1 μM). *n* = 8; 2-way ANOVA with Sidak’s multiple comparisons test. (Right) Fold change of isoproterenol-induced luciferase activity (isoprot/DMSO) in the presence of vehicle vs. indicated drugs (using the same data sets from the left graphs in E-G); two-tailed *t*-test. All data except in D: mean ± SD; data in D: median with interquartile range.

Finally, we examined if cytoplasmic Ca^2+^ increase is required for isoproterenol-induced ER-Mito coupling. A recently identified calcium mobilization inhibitor ebselen (30), eliminated the positive effect of isoproterenol (Fig. 5E), proving the requirement of cytoplasmic Ca^2+^ in isoproterenol-induced contacts. However, considering its multiple inhibitory functions, the possibility of ebselen inhibiting other factors or pathways could not be excluded. Unlike ebselen, the PLC inhibitors edelfosine and U73122 exhibited a rather mild inhibition on isoproterenol-induced ER-Mito contacts (Fig. 5F, G). When the fold changes of the isoproterenol-induced luciferase activity (over DMSO) were compared between the vehicle- vs. each drug-treated cells, edelfosine showed a trend toward blocking isoproterenol response (*p* = 0.0674) although the effects of both drugs did not reach statistical significance (Fig. 5F, G; right). These data argue for a significant involvement of Ca^2+^ but a negligible role of PLC in β-AR signaling-induced ER-Mito coupling.

### Actin polymerization is required for β-AR signal-mediated ER-Mito contact regulation

Finally, we explored the potential mechanism by which cytosolic Ca^2+^ and EPAC activation could modulate ER-Mito interaction. It has been shown that cytosolic Ca^2+^ spike by multiple stimuli such as ionomycin and histamine, triggers a transient surge in cytosolic actin polymerization (32, 33), which is required for ER-Mito contacts (28). Thus, we postulated that an actin polymerization ‘burst’ (achieved through cytosolic Ca^2+^ rise and/or other means upon β-AR activation) would be required for isoproterenol-induced ER-Mito interaction. To test this hypothesis, the cells were treated with latrunculin A, an actin polymerization inhibitor, and its effect on isoproterenol activity was assessed. Supporting our hypothesis, latrunculin abolished the isoproterenol-induced split-Rluc activity (Fig. 6B). This finding supports a model that actin filament assembly triggered by β-AR activation may facilitate a close contact between the ER and Mito.

**Figure 6.**
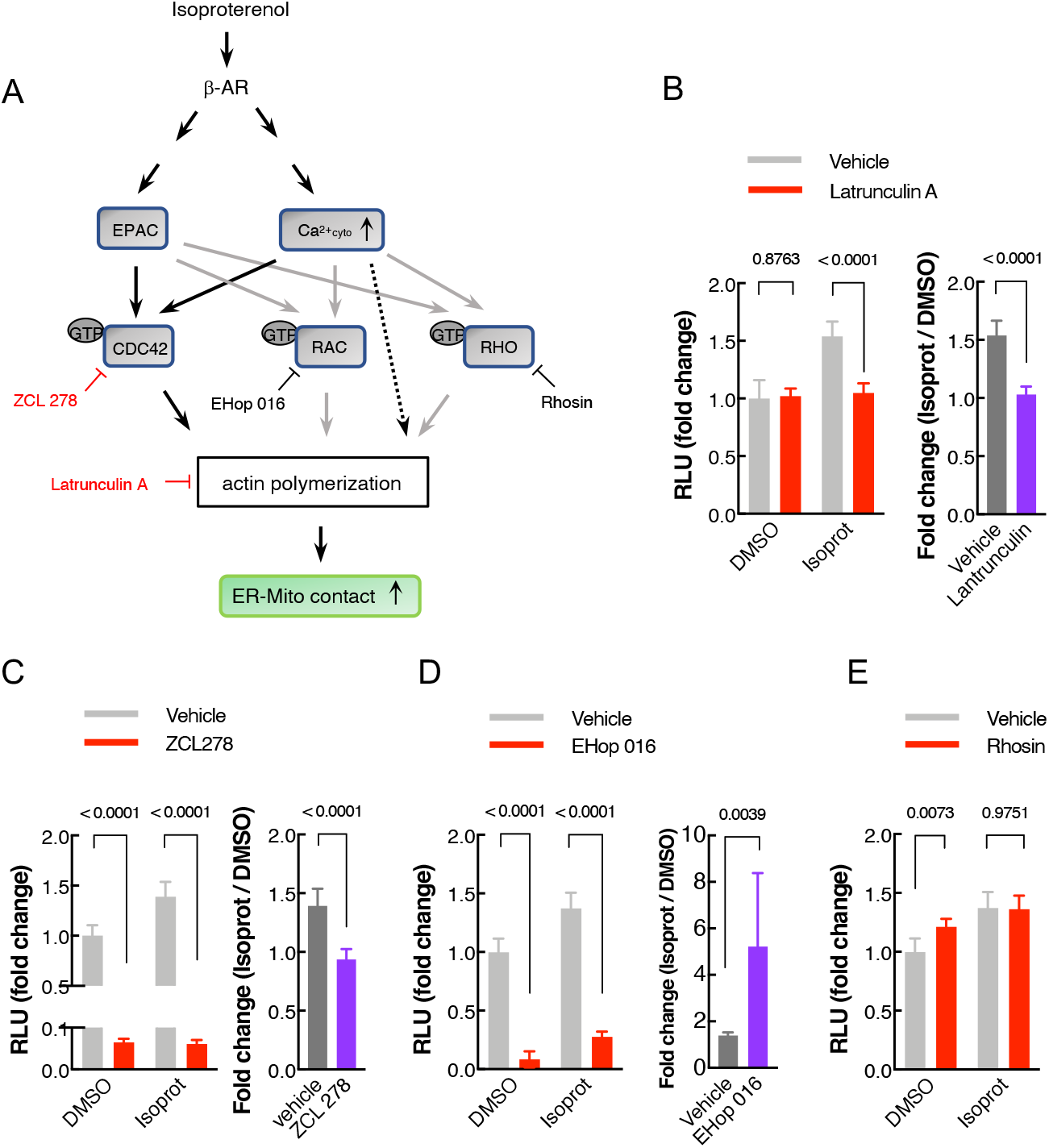
Actin polymerization is required for isoproterenol-induced ER-Mito contact regulation. ***A***, A schematic diagram (simplified) displaying the Rho GTPases and their upstream regulators potentially involved in actin polymerization. Arrow: pathways potentially involved (black) or not-involved (grey) in β-AR-mediated ER-Mito contact modulation. Drugs with red color: the ones blocking isoprot-induced split-Rluc activity; with black color, not blocking. ***B-D***, (Left): Split-Rluc assay in HEK293T cells treated with vehicle or indicated drugs (Latrunculin, 1 μM; ZCL 278, 25 μM; EHop 016, 12.5 μM) in the presence of DMSO or isoprot (1 μM). Fold changes: normalized to the mean of DMSO/vehicle. n = 8 (B), 16 (C), 8 (D); 2-way ANOVA with Sidak’s multiple comparisons test. (Right): Fold change of isoproterenol-induced luciferase activity (isoprot/DMSO) in the presence of vehicle vs. indicated drugs (same data from the left graphs in B-D). Two-tailed *t*-test. ***E***, Split-Rluc assay in HEK293T cells treated with vehicle or Rhosin (25 μM) in the presence of DMSO or isoprot (1 μM). Fold changes: normalized to the mean of DMSO/vehicle. *n* = 8; 2-way ANOVA with Sidak’s multiple comparisons test. All data: mean ± SD.

In the cell, actin reorganization is mainly regulated by RHO GTPases; CDC42, RAC, and RHO are the most prominent members among them (Fig. 6A). It has been well established that the activation of CDC42, RAC, and RHO promotes actin polymerization (Fig. 6A) (34). These RHO GTPases are regulated by GEFs (guanine nucleotide exchange factors), GAPs (GTPase activating proteins) and GDIs (guanine nucleotide dissociation inhibitors); the GPCR signaling is linked to the activation of these regulators (35, 36). In addition to G_12_-mediated pathway which is well documented for regulating RHO GTPases through RHO GEFs, increased intracellular Ca^2+^ (via activating calmodulin) as well as EPAC play roles in activating RHO GTPases (36–38). This prompted us to test if CDC42, RAC, and RHO participate in β-AR-induced ER-Mito contact formation.

To address this, CDC42, RAC, or RHO activity was pharmacologically inhibited in the presence of isoproterenol and then ER-Mito contacts were assayed. The CDC42 inhibitor ZCL 278 completely eradicated the effect of isoproterenol (Fig. 6C), supporting a role for CDC42 in β-AR-induced ER-Mito contacts, likely by triggering actin polymerization. This result was corroborated by PLA results where ZCL 278 counteracted the positive effect of forskolin on ER-Mito contacts (Fig. 4E). The RAC inhibitor EHop 016 also reduced split-Rluc activity when treated with or without isoproterenol (Fig. 6D, left). However, its inhibitory effect appeared to be independent of isoproterenol response, considering that the fold increase by isoproterenol over DMSO was in fact higher in EHop 016-treated cells when compared to vehicle-treated ones (Fig. 6D, right). Finally, when RHO activity was inhibited by rhosin, no changes in isoproterenol response was detected (Fig. 6E). These results strongly support that actin polymerization triggered by CDC42, but not by not RAC or RHO, is a downstream event facilitating ER-Mito coupling, in response to β-AR signal.

Of note, another cytoskeletal drug nocodazole, which interferes microtubule polymerization and blocks mitochondrial motility (39), caused a dose-dependent increase in split-Rluc activity; this suggests that microtubule polymerization normally hinders ER-Mito contacts probably due to excess mitochondrial dynamics along the microtubule (Fig. S3D; see Discussion). Together these results implicate an important role of cytoskeletal dynamics in ER-Mito contact modulation.

## Discussion

Communication between the ER and Mito via membrane contact sites is crucial for many cellular functions. Although recent studies have focused on identifying the molecular components of the contacts, the signaling pathway(s) involved in contact regulation remains unknown. Herein we have identified β-AR signaling as a significant pathway modulating ER-Mito interactions. Upon ligand binding, β-ARs transduce the signal through G_s_/AC/cAMP/EPAC and Ca^2+^ to induce physical contacts between the ER and Mito likely by stimulating a rapid, transient actin filament assembly through CDC42 activation. As a result, mitochondrial Ca^2+^ uptake is enhanced, thus activating mitochondrial matrix proteins to generate more ATPs (Fig. 7) (20, 23). Together our data provide significant insights into the roles of ER-Mito contacts in maintaining cellular homeostasis and in responding to physiological demands or stresses, under the influence of the GPCR signaling.

**Figure 7.**
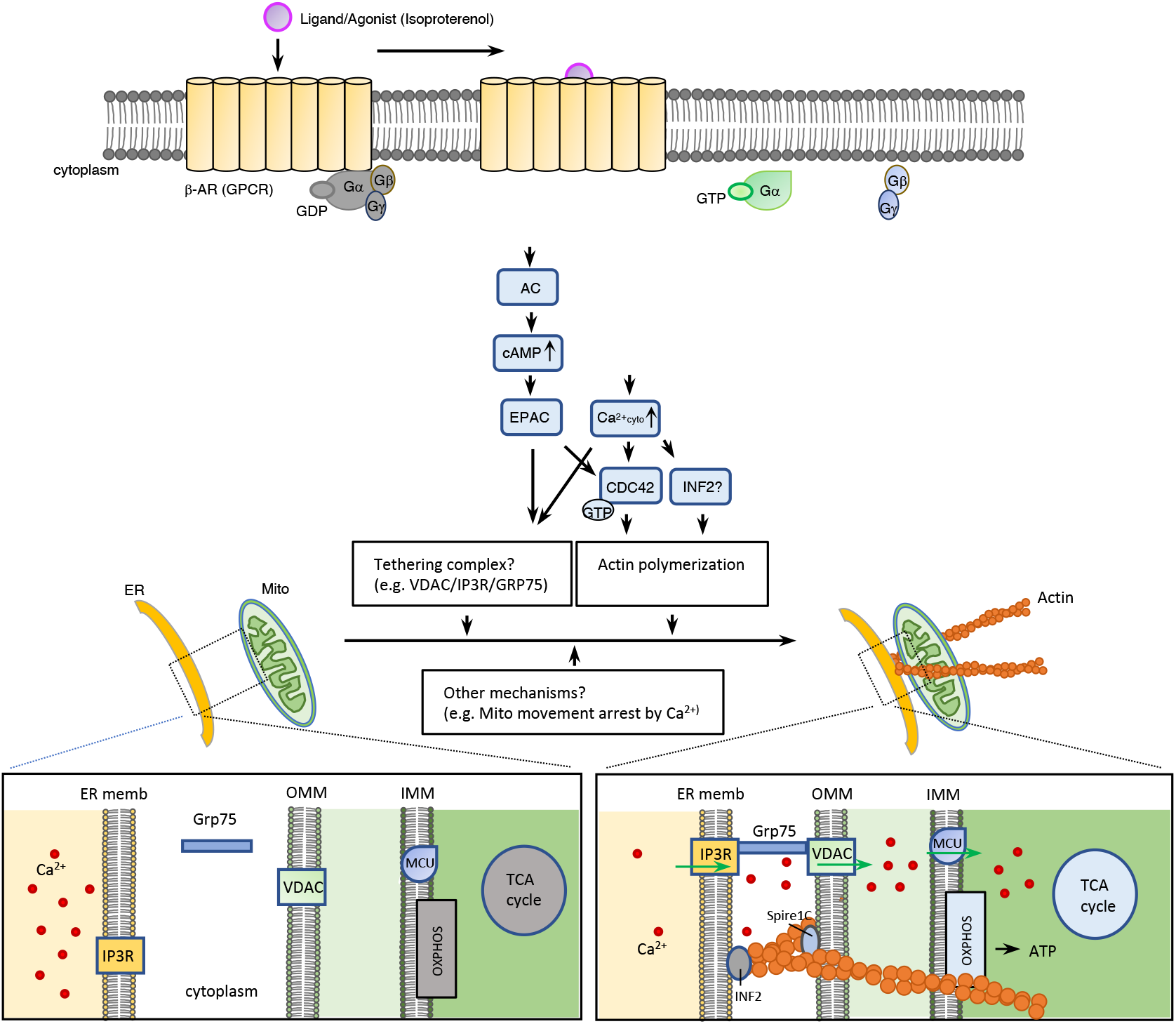
Proposed model for potential mechanisms of β-AR (GPCR) signaling in ER-Mito contact regulation. We postulate that the increase in cAMP and subsequent activation of EPAC facilitates ER-Mito interactions likely through 1) triggering actin polymerization by CDC42; 2) inducing tethering complex formation (e.g., VDAC1/IP3R1/GRP75). Increased cytosolic Ca^2+^ can also increase ER-Mito interactions by influencing actin polymerization through CDC 42 or potentially through INF2 (inverted formin2) (ref. 31). It is also plausible that cAMP and Ca^2+^ might employ other mechanisms (e.g., Mito movement arrest on microtubule by Ca^2+^) and that another downstream pathway (independent of Gs or Gq) of β-AR might also participate in ER-Mito coupling. β-AR-mediated ER-Mito coupling increases mitochondrial Ca^2+^ uptake, which enhances mitochondrial bioenergetics by activating proteins involved in TCA cycle and oxidative phosphorylation (OXPHOS).

Our data support a model where EPAC activation via G_s_/AC/cAMP, and cytosolic Ca^2+^ increase leads to CDC42 activation (36–38) and subsequently to actin filament assembly (34), a driving force for ER-Mito interaction (Fig. 7). In addition to CDC42, ER-bound inverted formin2 (INF2), could be a good candidate protein modulating actin filament assembly, given that it mediates actin polymerization in response to cytosolic Ca^2+^ burst (28) upon GPCR activation (40). Furthermore, INF2-mediated actin polymerization at the ER increases ER-Mito contacts, Mito Ca^2+^ uptake, and mitochondrial fission (28), all of which were also increased with isoproterenol treatment (Fig. 2, 3).

In addition to stimulating actin dynamics, EPAC and cytosolic Ca^2+^ surge may also exert their effects through other mechanisms. For instance, they might facilitate the formation of ER-Mito coupling complexes. In fact, a recent study demonstrated that under stress condition, EPAC enhances ER-Mito coupling by promoting the interaction between VDAC1/IP3R1/GRP75, an ER-Mito tethering complex (27). In a similar way, Ca^2+^ might affect the tethering by controlling either the tethering proteins directly or modulators of the complex. For example, the presence of Ca^2+^-responsive tethering proteins such as E-Syt1 (extended synaptotagmin 1), at the ER-plasma contact sites (41), raises the possibility of the existence of similar proteins at the ER-Mito contact sites. In this regard, it will be interesting to investigate whether IP3Rs (IP3R1, 2, and 3) represent such tethering proteins responding to cytoplasmic Ca^2+^ surge, given their structural roles independent of their channel functions (42) and the presence of Ca^2+^ binding sites on them (43). Another possible mechanism by which cytosolic Ca^2+^ regulates ER-Mito coupling, is through modulating mitochondrial dynamics. It has been documented that IP3R-mediated cytosolic Ca^2+^ rise arrests mitochondrial movement along the microtubule, increasing the likelihood of its interaction with ER (39). Consistent with this, nocodazole (an inhibitor of microtubule polymerization shown to block mitochondrial motility) increased ER-Mito contacts in a dose dependent manner (Fig. S3D).

Accumulating evidence indicates that ER-Mito dysfunction contributes to a variety of neurodegenerative diseases (4, 44). In Parkinson’s disease models, β2-AR agonists have been documented to be beneficial (45). Salbutamol and metaproterenol (β2-AR agonists), both of which were also identified in our screen, reduce α-syn transcription levels and appears to reduce risk of developing PD (14, 15). Although their efficacy was linked to α-syn transcription regulation (14) and anti-inflammation (46), our data indicate that these drugs might also exert therapeutic effects by increasing ER-Mito contacts since some PD-associated mutation insults disrupt these contacts (47). This opens up the possibilities for therapy of other conditions associated with ER-Mito contact dysregulation. When considering ER-Mito contact enhancement drugs for therapeutic options, it is important to recognize that both increased and decreased contacts can be detrimental. Therefore, careful titration of these drugs will be crucial.

Our results demonstrate that the split-Rluc assay can serve as a valid assay to screen for modulators of ER-Mito contacts, yet we recognize its limitation. Given that it takes approximately 1.5 hr for EnduRen substrate to generate stable luminescence, which then will last for 24 hours, the split-Rluc assay is not suitable to detect real-time dynamics of ER-Mito contacts. This explains the discrepancy between the PLA and split-Rluc assay when comparing the signal intensities in time course experiments for isoproterenol treatment; PLA signal shows increase (up to 3 hr) then decrease (at 6hr) (Fig. S2B) probably due to a negative feedback in GPCR signaling, while the split-Rluc activity shows continuous increase up to 16 hrs (maximum time measured) (data not shown). The split-Rluc assay appears to detect accumulated changes (irreversible), despite the relatively short half-life of each split fragment (Fig. S1F).

While our current screen focused on an FDA-approved drug library, further screens with other compound libraries will likely provide additional biologic insights into this important inter-organelle communication (e.g., identifying small molecules that could directly target ER-Mito contact sites). Furthermore, ORF or siRNA library screens would be warranted to identify additional molecular machinery functioning for the contact modulation. Finally, while our current screening system is most suitable for enhancer identification due to low basal level of split-Rluc activity, modification to our system could permit the screening for inhibitors in future studies. Furthermore, the detailed mechanisms of the GPCR pathway-regulated ER-Mito contacts and any distinctions, among different classes of GPCRs, in downstream pathways leading to ER-Mito contact modulation will need to be addressed.

## Materials and Methods

### Plasmids and DNA constructs

Split-Rluc constructs (Mito-Rluc^N^ and Rluc^C^-ER) have been described previously (8). R-GECO1 (plasmid #46021), GCaMP6s (plasmid #100844), and Sec61β (plasmid #15108) were obtained from Addgene (gift from Robert Campbell, Douglas Kim & GENIE Project, and Tom Rapoport, respectively). CRE-luc2P (pGL4.29) used for CRE reporter gene assay was purchased from Promega. Plasmids expressing Gs CA (Q227L), Gq CA (Q209L), Gi CA (Q205L), RAP1A DN (S17N), and EPAC2 DN (R432K) (48–51) were generated by cloning PCR-amplified cDNA fragments into pCAG-IRES-GFP (pCIG) vector (EcoRI/MluI). For the plasmid encoding β2-AR was generated by cloning PCR-amplified cDNA fragment into pCDH-EF1a-MCS-PGK-puro vector (EcoRI/NotI). For these cloning, HEK293T cDNAs, as a template, and specific primer sets including the ones containing mutations (see Table S5 for primer sequences) were used for PCR.

### Compounds and chemical reagents

We used NIH Clinical Collections 1 and 2 (NCC) (Evotec, San Francisco, CA) containing 720 FDA approved drugs dissolved in 100% DMSO at a concentration of 10 mM (the original list contains 727 drugs; 7 of them duplicated). All drugs were used at 1 μM final concentration. The target pathway(s) of the members of this collection was manually curated from the information available in the PubChem portal (http://pubchem.ncbi.nlm.nih.gov/). For post-screen experiments, the chemical reagents were used as following: isoproterenol (1747, Tocris, 1μM), sotalol (0952, Tocris, 250 μM), actinomycin D (A9415, Sigma-Aldrich, 0.8 μM), cycloheximide (asd, 20 μg/ml), CCPA (119136, Sigma-Aldrich, 1μM), prostaglandin E1 (P5515, Sigma-Aldrich, 1μM), ESI 09 (4773, Tocris, 5 μM), cAMPS-Rp (13371, Tocris, 10 μM), H89 (2910, Tocris, 10 μM), PD0325901 (04-0006, Reprocell, 1 μM), histamine (H7250, Sigma-Aldrich, 100 μM), ionomycin (1704, Tocris, 1 μM), and cyclopiazonic acid (CPA) (C1530, Sigma-Aldrich, 10 μM), thapsigargin (T9033, Sigma-Aldrich, 0.1-10 μM), ebselen (5245, Tocris,, 50 μM), edelfosine (3022, Tocris, 40 μM), U73122 (1268, Tocris, 5 μM), latrunculin (ab144290, Abcam, 1 μM), ZCL278 (4794, Tocris, 25 μM), EHop016 (6248, Tocris, 12.5 μM), rhosin (5003, Tocris, 25 μM), and nocodazole (S2775, Selleck Chemicals, 1, 4, and 10 ng/ml). For DMSO used as controls, the final concentration was decided according to the experimental drug’s final concentration (e.g., 0.01% DMSO was used as a control, when other drugs (in 100 % DMSO) were used 1:10000 dilution).

### Cell culture and transfection

HEK293T, HeLa, C2C12, and Neuro2a cells were maintained in DMEM (Thermo Fisher Scientific) with 10% (HEK293T, HeLa, Neuro2a) or 15% (C2C12) FBS (Thermo Fisher Scientific) in a humidified incubator at 37°C with 5% CO_2_. For transfection in 8-well chamber slides, 12-well plates, and 100 mm culture dishes, cells were plated at a density of 3 × 10^5^/well, 4 × 10^5^ /well, and 5 × 10^6^/well respectively to obtain 75-80% confluency, and 100 ng, 400 ng, and 5 μg each of DNA constructs respectively were used with polyethylenimine (PEI; Polysciences, Eppelheim, Germany) as previously described (8, 16). Of note, we have used different batches of HEK293T cells for the initial screen vs the subsequent experiments.

### Split-Renilla luciferase (split-Rluc) reconstitution assay

HEK293T cells were transfected with split-Rluc reporter plasmids (Mito-Rluc^N^ and Rluc^C^-ER) in 12 well plates. Four to six hours after transfection, cells were dissociated and re-plated into a 96-well plate coated with poly-D-lysine (50 μg/mL). Twenty-four hours post-transfection, live cell substrate EnduRen (30 μM, Promega) was added to the culture media and incubated for 2-5 hours. Luciferase activity (luminescence) was measured by a POLARstar Omega microplate reader (BMG LABTECH, Ortenberg, Germany) and normalized to the mean of the control.

### Library screen

HEK293T cells were transfected with split-Rluc constructs in a 100 mm dish and used for split-Rluc assay. 1 μM of each compound (final concentration) was added to each well together with Enduren and incubated for 5 hours followed by luminescence measurement. The initial screen was carried out once. The split-Rluc activity of each drug was normalized by quantile normalization using R software (version 4.0). Hits were picked using the cut-off value of 3SD higher than the average of the control (DMSO), and further tested for their activity with one more round of split-Rluc assay. For drug enrichment analysis, we employed the over representation analysis (52). False discovery rate (FDR) was calculated by R software. All dendrograms were created using R with packages ‘ggraph’, ‘igraph, ‘tidyverse’ and ‘RcolorBrewer’.

### Proximity labeling assay (PLA)

HeLa cells were transfected with SEC61-GFP expression vector in 8 well chamber slide (Millipore Sigma, Millicell EZ SLIDE) and, 16 hours after transfection, treated with DMSO or indicated drugs for 5 hrs. PLA was performed according to the manufacturer’s recommendation (Duolink^™^ In Situ Red Starter Kit Mouse/Rabbit, Sigma-Aldrich). Primary antibodies against GFP (ab13970, chicken polyclonal, Abcam) and Tom20 (sc-17764, mouse monoclonal, Santa Cruz Biotechnology) were used at 1:500 dilution each. After mounting, the images were captured by Zeiss Observer Z1 inverted microscope (Hamamatsu ORCA-Flash4.0 camera) using Zeiss Zen Pro software and analyzed in ImageJ with Maxima function. For analyses, all PLA studies were internally compared.

### Transmission Electron Microscopy (TEM)

HEK293T cells cultured in 100 mm dish were treated with DMSO or isoproterenol for 30 min. These monolayers of cells were fixed with glutaraldehyde/paraformaldehyde mixture and post-fixed in 2% aqueous OsO_4_ in 0.2M S-collidine buffer, pH 7.4. One-micron thick sections were cut on an ultra-microtome and stained on the glass slide, and thin sections were mounted on 200 mesh copper grids and stained as previously described (53, 54). The stained grids were examined on a Jeol JEM-100CX electron microscope equipped with a digital camera. With randomly selected EM images (160 images for control and 158 images for isoproterenol), the distance between the outer mitochondrial membrane (OMM) and ER membrane was measured. If the distance (measured in Image J) was within 30 nm, it was counted as ‘contact’, whereas if greater than 30 nm, ‘no contact’. The number of mitochondria with ER contact was counted and its percentage over the total number of mitochondria was calculated. For the total contact coverage, the partial mitochondrial perimeter showing a close contact with ER was measured and added all together. This sum was divided by the sum of the mitochondrial perimeter measured among all mitochondria regardless of their contact status. For the contact length per mitochondrion, the partial perimeter of each mitochondrion showing contact with ER membrane (less than 30 nm distance) and the entire perimeter were measured for each mitochondrion in ImageJ and the ratio was plotted.

### Mito-APEX proximity labeling

HEK293T cells were transfected with pcDNA3-mito-APEX in a 6-well plate as described in supplementary Methods. For proximity labeling of proteins, previously reported method (16) was used with a slight modification. Briefly, cells were treated with DMSO or isoproterenol (1 μM) for 1 hr, before incubated with biotin-phenol (500 μM) and treated with H_2_O_2_ for 3 min (16). After the biotinylation reaction was quenched (16), cells were lysed with RIPA buffer and centrifuged at 13,000 rpm for 10 min at 4°C. To get rid of the free biotin-phenol, the sample was applied to Zeba Spin Desalting Column (Thermo Fisher Scientific). The recovered lysates were incubated with streptavidin magnetic beads (16), and the beads were washed five times with RIPA buffer. Biotinylated proteins of mito-APEX pull downs were eluted by boiling with 2X protein loading buffer (Bio-Rad) containing 2 mM biotin and 20 mM DTT at 95 °C for 10 min and subjected to 4-15% SDS-PAGE followed by Western blot analysis described below.

### Western blot analysis

For Western blots of affinity purified biotinylated proteins by Mito-APEX, we followed previously described procedures (16). The following primary antibodies were used: IP3R antibody (PA1-901, Thermo Scientific, rabbit polyclonal, 1:1000), GFP antibody (sc-8334, Santa Cruz Biotechnology, rabbit polyclonal, 1:500), and GAPDH antibody (CB1001, Calbiochem, mouse monoclonal, 1:5000). For Western blots of Rluc^C^-ER and Mito-Rluc^N^, HEK293T cells were transfected with split-Rluc constructs as described above. Cells treated with DMSO, isoproterenol, actinomycin, or isoproterenol/actinomycin for 1 hr were lysed and subjected to 4-15% SDS-PAGE followed by Western blotting as described before (16). Primary antibodies against MYC (2276, 9E10, Cell signaling, mouse monoclonal, 1:1000), FLAG (F1804, M2, Sigma, mouse monoclonal, 1:1000), and actin (1844-1, Epitomics, rabbit monoclonal, 1:2000), and the appropriate secondary antibodies were used. For protein half-life analysis, HEK293T cells were treated with actinomycin D and cycloheximide together and harvested at different time points as indicated.

### Calcium level measurement

To monitor the mitochondrial and cytoplasmic calcium dynamics, mito-R-GECO1 and GCaMP6s constructs respectively were transfected into HeLa cells in an 8-well chamber slide. Sixteen hours after transfection, the cells were treated with DMSO or indicated drugs and imaged on Zeiss Observer Z1 inverted microscope using Zeiss Zen Pro software, every 2.5 or 5 sec. Approximately 7.5 sec (histamine) or 15 sec (ionomycin) after the start of the experiment, histamine (100 μM) or ionomycin (3 μM) was superfused over transfected cells. The boundary of each cell was set as ROIs (regions of interest) and randomly selected based on the presence of R-GECO1 or GCaMP6s fluorescent signal. ΔF/F_0_ and (ΔF/F_0_)/Δt (R-GECO1) (from 7.5 to 12.5 sec) were calculated as described previously (55).

### Mitochondrial transmembrane potential and mitochondrial length

Mitochondrial membrane potential (Ψ_mit_) was measured by tetramethyl rhodamine methyl ester (TMRM) fluorescent dye (I34361, Thermo Fisher Scientific). HeLa cells were loaded with 10 nM TMRM for 30 min at RT and then treated with DMSO or isoproterenol (1 μM) for 1 hour. TMRM fluorescence was imaged using Zeiss Observer Z1 inverted microscope, and the signal intensity was analyzed in Image J. The lengths of mitochondria were analyzed in Image Analyst MKII software (version 4.0) with default parameter [largest mitochondria size (width in pixel) = 5, Sensitivity (top range scaling, percentile) = 99.5 and minimum size (area, pixels) = 5].

### GPCR expression level analysis using RNA-seq data

The raw sequence files were downloaded from NIH GEO data sets with each GEO set access number (GSE99249 for HEK293T, GSE122986;SRR8247560 for HeLa, GSE148617;SRR10125054 for C2C12, GSE84481;SRR3927297 for Neuro2a). The downloaded fastq files were aligned and sorted with GRCh38 or GRCm38 genome index by HISAT2 (version 2.1.0) and samtools (version 1.9), respectively. We calculated TPM with TPMCalculator (56).

### Statistical analysis

All statistical analyses were performed using GraphPad Prism (version 8.4.2 or 9.1.0) or R (version 4.0.0). To determine whether differences between two data sets are statistically significant, a two-tailed *t*-test was used if the data passed normality test; a non-parametric test (Mann Whitney test) was used if failed normality test. When more than two data sets were compared, a 1- or 2-way ANOVA test was used if passed normality test; other tests recommended by Prism software (e.g., Kruskal-Wallis test) were used if data failed the normality test. For comparisons of multiple samples following these tests, Dunn’s test, Dunnett’s test, or Sidak’s test was used as recommended by Prism. For data normality analysis, Anderson-Darling, D’Agostino-Pearson, Shapiro-Wilk, and Kolmogorov-Smirnov tests were used. For the categorical data comparison, a Chi-square test was used. The details of all statistical tests including the exact *p*-values and sample numbers (*n*) are listed in the figure legends or directly in the graphs. All samples (e.g., cells, mitochondria, and ROIs etc.) were randomly selected, and image quantifications were carried out blindly. *P* values ≤ 0.05 were considered statistically significant. When fold changes were used, all data were normalized to control (e.g., DMSO). All bar graphs are plotted as mean ± standard deviation (SD). Concentration responsive curves are plotted as mean ± SEM. All scattered plots are marked with individual data points and median with inter quartile range.

## Acknowledgments

This work was supported by NINDS (R01NS100007) and by Dana-Farber Cancer Institute microgrant (#120205). The authors are grateful to Dr Jeffrey A. Golden and his lab members (BWH/HMS) for their support and critical review of the manuscript, to Dr Sandro Santagata (BWH/HMS) for a critical review of the manuscript, to Drs Sean E. Lawler and Michal Oscar Nowicki (BWH/HMS) for sharing their reagents and equipment, to Amanda Frischmann for her assistance on compound list curating process, and to Dr Wei Wang (BWH/HMS) for her help with statistical analyses supported by the Harvard Catalyst|The Harvard Clinical and Translational Science Center (NIH award UL 1TR002541). The content is solely the responsibility of the authors and does not necessarily represent the official views of Harvard Catalyst, Harvard University and its affiliated academic healthcare centers, or the NIH.

## References

1. Csordás G, Weaver D, Hajnóczky G (2018) Endoplasmic Reticulum-Mitochondrial Contactology: Structure and Signaling Functions. Trends in Cell Biology 28(7):523–540.

2. Kornmann B, et al. (2009) An ER-mitochondria tethering complex revealed by a synthetic biology screen. Science 325(5939):477–481.

3. Eisenberg-Bord M, Shai N, Schuldiner M, Bohnert M (2016) A Tether Is a Tether Is a Tether: Tethering at Membrane Contact Sites. Dev Cell 39(4):395–409.

4. Paillusson S, et al. (2016) There’s Something Wrong with my MAM; the ER-Mitochondria Axis and Neurodegenerative Diseases. Trends Neurosci 39(3):146–157.

5. Moltedo O, Remondelli P, Amodio G (2019) The Mitochondria-Endoplasmic Reticulum Contacts and Their Critical Role in Aging and Age-Associated Diseases. Front Cell Dev Biol 7:172.

6. Rieusset J (2018) The role of endoplasmic reticulum-mitochondria contact sites in the control of glucose homeostasis: an update. Cell Death and Disease 9(3):388.

7. Doghman Bouguerra M, Lalli E (2019) ER-mitochondria interactions: Both strength and weakness within cancer cells. Biochim Biophys Acta Mol Cell Res 1866(4):650–662.

8. Lim Y, Cho I-T, Schoel LJ, Cho G, Golden JA (2015) Hereditary spastic paraplegia-linked REEP1 modulates endoplasmic reticulum/mitochondria contacts. Ann Neurol 78(5):679–696.

9. Tepikin AV (2018) Mitochondrial junctions with cellular organelles: Ca2+ signalling perspective. Pflugers Arch - Eur J Physiol 470(8):1181–1192.

10. Le Masson G, Przedborski S, Abbott LF (2014) A computational model of motor neuron degeneration. Neuron 83(4):975–988.

11. Vassilatis DK, et al. (2003) The G protein-coupled receptor repertoires of human and mouse. Proc Natl Acad Sci USA 100(8):4903–4908.

12. Weis WI, Kobilka BK (2018) The Molecular Basis of G Protein-Coupled Receptor Activation. Annu Rev Biochem 87:897–919.

13. Inoue A, et al. (2019) Illuminating G-Protein-Coupling Selectivity of GPCRs. Cell 177(7):1933–1947.e25.

14. Mittal S, et al. (2017) β2-Adrenoreceptor is a regulator of the α-synuclein gene driving risk of Parkinson’s disease. Science 357(6354):891–898.

15. Gronich N, et al. (2018) β2-adrenoceptor agonists and antagonists and risk of Parkinson’s disease. Mov Disord 33(9):1465–1471.

16. Cho I-T, et al. (2017) Ascorbate peroxidase proximity labeling coupled with biochemical fractionation identifies promoters of endoplasmic reticulum-mitochondrial contacts. J Biol Chem 292(39):16382–16392.

17. Söderberg O, et al. (2006) Direct observation of individual endogenous protein complexes in situ by proximity ligation. Nat Meth 3(12):995–1000.

18. Zhao Y, et al. (2011) An expanded palette of genetically encoded Ca^2+^ indicators. Science 333(6051):1888–1891.

19. Filadi R, et al. (2018) TOM70 Sustains Cell Bioenergetics by Promoting IP3R3-Mediated ER to Mitochondria Ca2+ Transfer. Curr Biol 28(3):369–382.e6.

20. Bravo R, et al. (2011) Increased ER-mitochondrial coupling promotes mitochondrial respiration and bioenergetics during early phases of ER stress. J Cell Sci 124(Pt 13):2143–2152.

21. Cárdenas C, et al. (2010) Essential regulation of cell bioenergetics by constitutive InsP3 receptor Ca2+ transfer to mitochondria. Cell 142(2):270–283.

22. Territo PR, Mootha VK, French SA, Balaban RS (2000) Ca(2+) activation of heart mitochondrial oxidative phosphorylation: role of the F(0)/F(1)-ATPase. Am J Physiol, Cell Physiol 278(2):C423–35.

23. Denton RM (2009) Regulation of mitochondrial dehydrogenases by calcium ions. Biochim Biophys Acta 1787(11):1309–1316.

24. Perry CGR, Kane DA, Lanza IR, Neufer PD (2013) Methods for assessing mitochondrial function in diabetes. Diabetes 62(4):1041–1053.

25. Friedman JR, et al. (2011) ER tubules mark sites of mitochondrial division. Science 334(6054):358–362.

26. Huang X-P, Song X, Wang H-Y, Malbon CC (2002) Targeted expression of activated Q227L G(alpha)(s) in vivo. Am J Physiol, Cell Physiol 283(2):C386–95.

27. Fazal L, et al. (2017) Multifunctional Mitochondrial Epac1 Controls Myocardial Cell Death. Circulation Research 120(4):645–657.

28. Galaz-Montoya M, et al. (2017) β2-Adrenergic receptor activation mobilizes intracellular calcium via a non-canonical cAMP-independent signaling pathway. J Biol Chem 292(24): 9967–9974.

29. Chakrabarti R, et al. (2018) INF2-mediated actin polymerization at the ER stimulates mitochondrial calcium uptake, inner membrane constriction, and division. J Cell Biol 217(1):251–268.

30. Plenge-Tellechea F, Soler F, Fernandez-Belda F (1997) On the inhibition mechanism of sarcoplasmic or endoplasmic reticulum Ca2+-ATPases by cyclopiazonic acid. J Biol Chem 272(5):2794–2800.

31. Kuschak M, et al. (2020) Cell-permeable high-affinity tracers for Gq proteins provide structural insights, reveal distinct binding kinetics and identify small molecule inhibitors. British Journal of Pharmacology 177(8):1898–1916.

32. Ji W-K, et al. (2017) Receptor-mediated Drp1 oligomerization on endoplasmic reticulum. J Cell Biol 216(12):4123–4139.

33. Shao X, Li Q, Mogilner A, Bershadsky AD, Shivashankar GV (2015) Mechanical stimulation induces formin-dependent assembly of a perinuclear actin rim. Proceedings of the National Academy of Sciences 112(20):E2595–601.

34. Hall A (1998) Rho GTPases and the actin cytoskeleton. Science 279(5350):509–514.

35. Sinha S, Yang W (2008) Cellular signaling for activation of Rho GTPase Cdc42. Cell Signal 20(11):1927–1934.

36. Momotani K, Somlyo AV (2012) p63RhoGEF: a new switch for G(q)-mediated activation of smooth muscle. Trends Cardiovasc Med 22(5):122–127.

37. van Unen J, et al. (2016) Kinetics of recruitment and allosteric activation of ARHGEF25 isoforms by the heterotrimeric G-protein Gαq. Scientific Reports 6(1):36825.

38. Jin M, et al. (2005) Ca2+-dependent regulation of rho GTPases triggers turning of nerve growth cones. J Neurosci 25(9):2338–2347.

39. Yi M, Weaver D, Hajnóczky G (2004) Control of mitochondrial motility and distribution by the calcium signal: a homeostatic circuit. J Cell Biol 167(4):661–672.

40. Wang Y, et al. (2019) GPCR-induced calcium transients trigger nuclear actin assembly for chromatin dynamics. Nature Communications:1–9.

41. Giordano F, et al. (2013) PI(4,5)P(2)-dependent and Ca(2+)-regulated ER-PM interactions mediated by the extended synaptotagmins. Cell 153(7):1494–1509.

42. Bartok A, et al. (2019) IP3 receptor isoforms differently regulate ER-mitochondrial contacts and local calcium transfer. Nature Communications 10(1):3726.

43. Taylor CW, Tovey SC (2010) IP(3) receptors: toward understanding their activation. Cold Spring Harbor Perspectives in Biology 2(12):a004010.

44. Wilson EL, Metzakopian E (2020) ER-mitochondria contact sites in neurodegeneration: genetic screening approaches to investigate novel disease mechanisms. Cell Death Differ. doi:10.1038/s41418-020-00705-8.

45. Peterson L, Ismond KP, Chapman E, Flood P (2014) Potential benefits of therapeutic use of β2-adrenergic receptor agonists in neuroprotection and Parkinsonμs disease. J Immunol Res 2014:103780.

46. Qian L, et al. (2011) β2-adrenergic receptor activation prevents rodent dopaminergic neurotoxicity by inhibiting microglia via a novel signaling pathway. J Immunol 186(7):4443–4454.

47. Barazzuol L, Giamogante F, Brini M, Calì T (2020) PINK1/Parkin Mediated Mitophagy, Ca2+ Signalling, and ER-Mitochondria Contacts in Parkinson’s Disease. IJMS 21(5). doi:10.3390/ijms21051772.

48. Huang X-P, Song X, Wang H-Y, Malbon CC (2002) Targeted expression of activated Q227L G(alpha)(s) in vivo. Am J Physiol, Cell Physiol 283(2):C386–95.

49. Murga C, Laguinge L, Wetzker R, Cuadrado A, Gutkind JS (1998) Activation of Akt/protein kinase B by G protein-coupled receptors. A role for alpha and beta gamma subunits of heterotrimeric G proteins acting through phosphatidylinositol-3-OH kinasegamma. J Biol Chem 273(30):19080–19085.

50. Dupuy AG, et al. (2020) Novel Rap1 dominant-negative mutants interfere selectively with C3G and Epac. Oncogene 24:4509–4520.

51. López De Jesús M, et al. (2006) Cyclic AMP-dependent and Epac-mediated activation of R-Ras by G protein-coupled receptors leads to phospholipase D stimulation. J Biol Chem 281(31):21837–21847.

52. Boyle EI, et al. (2004) GO::TermFinder--open source software for accessing Gene Ontology information and finding significantly enriched Gene Ontology terms associated with a list of genes. Bioinformatics 20(18):3710–3715.

53. Venable jh, Coggeshall R (1965) A simplified lead citrate stain for use in electron microscopy. J Cell Biol 25(2):407–408.

54. Reynolds ES (1963) The use of lead citrate at high pH as an electron-opaque stain in electron microscopy. J Cell Biol 17(1):208–212.

55. Fülöp L, Szanda G, Enyedi B, Várnai P, Spät A (2011) The effect of OPA1 on mitochondrial Ca^2+^ signaling. PLoS ONE 6(9):e25199.

56. Vera Alvarez R, Pongor LS, Mariño-Ramírez L, Landsman D (2019) TPMCalculator: one-step software to quantify mRNA abundance of genomic features. Bioinformatics 35(11):1960–1962.

