## Supplemental Figures and Tables for "G protein-coupled receptor signaling regulates ER-mitochondria contacts"

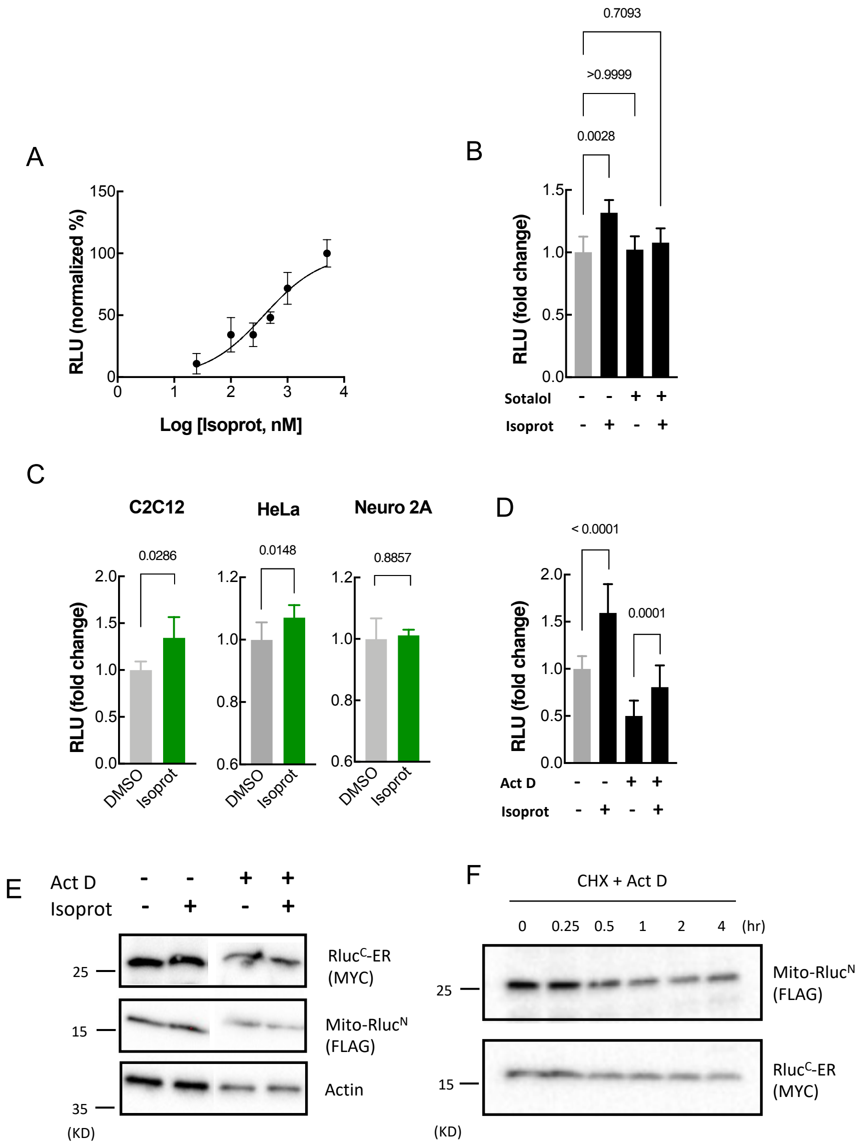


**Figure S1**. ***A,*** A concentration-responsive curve of isoproterenol effect on split-Rluc activity (*n* = 16; nonlinear regression (non-linear fit, EC50 = 385.4 nM); Error bars: SEM. ***B,*** The effect of β-AR antagonist (sotalol, 250 μM) on isoproterenol (isoprot)-induced split-Rluc activity. *n = 6*; Kruskal-Wallis test with Dunn's multiple comparisons test. All data: mean ± SD. ***C,*** Split-Rluc activities (RLU, relative light unit; normalized with the mean of DMSO) of HeLa, C2C12 or Neuro2a cells, treated with DMSO (control) or isoproterenol (1 μM). *n* = 4 (C2C12 and N2a), 8 (HeLa); Mann-Whitney test; All data: mean ± SD. ***D,*** The effect of the transcription inhibitor actinomycin D (ActD, 0.8 μM) on isoproterenol-induced split-Rluc activity. *n* = 15; 1-way ANOVA with Sidak's multiple comparisons test. All data: mean ± SD. ***E,*** The effect of Act D and isoproterenol on the expression level of split-Rluc fragments. Western blot analysis of split-Rluc fragments in HEK293T cells treated with indicated drugs, probed with MYC (for Rluc$^{C}$-ER), FLAG (for Mito-Rluc$^{N}$), or actin (loading control) antibody. ***F,*** Western blot analysis of protein levels for split-Rluc fragments in HEK293T cells treated with Act D (0.8 μM) and cycloheximide (protein translation inhibitor, 20 μg/ml). Samples were taken at the time points indicated and probed with MYC or FLAG antibody. *p* values (if ≤ 0.05, significant) are indicated at the top of each graph.

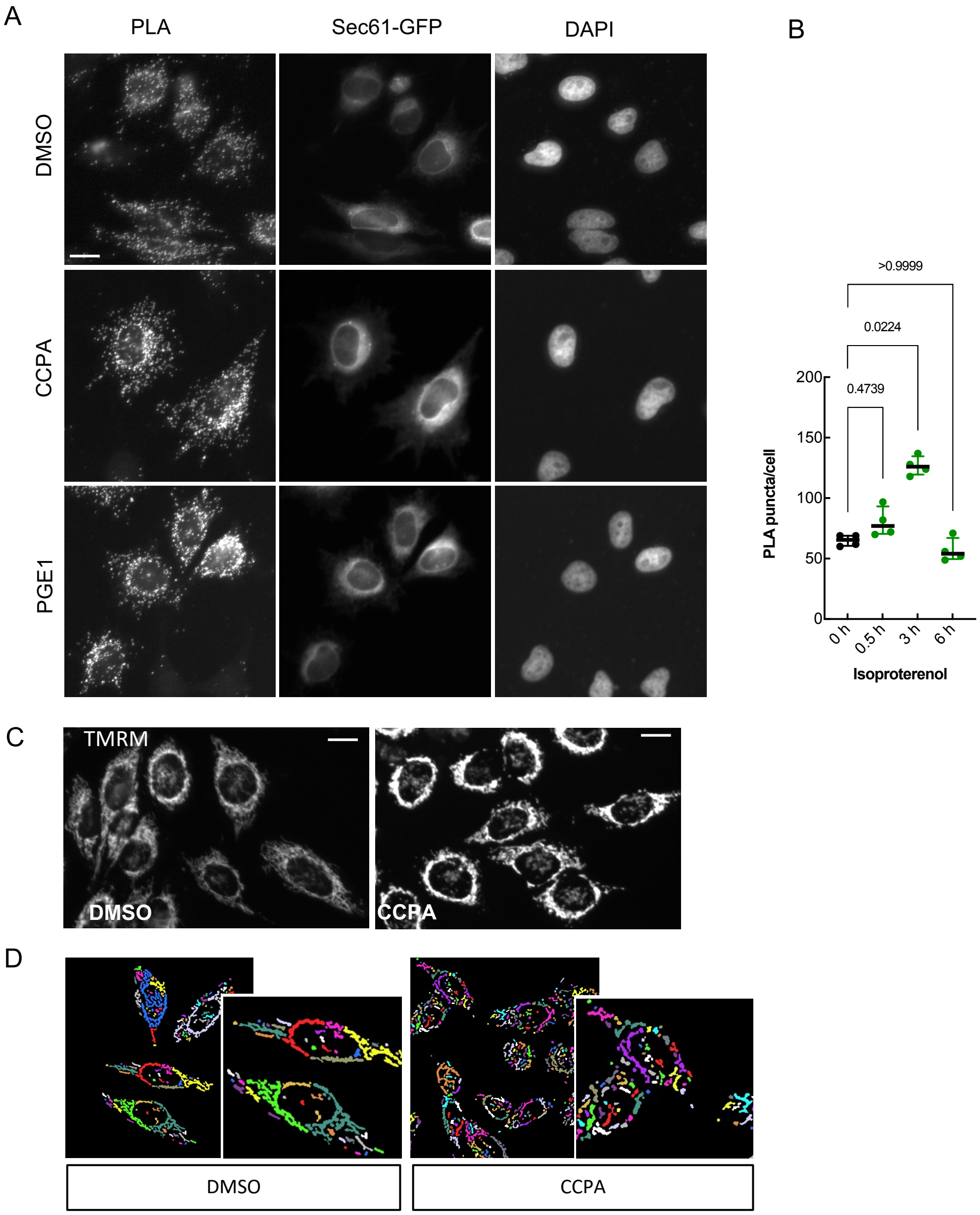


**Figure S2.** ***A,*** Representative images of proximity ligation assay (PLA) in HeLa cells treated with DMSO, CCPA, or PGE1 (1 μM each). PLA fluorescent signal indicates a close apposition between ER and Mito. Sec61-GFP, to label ER; DAPI, nucleus. Scale bars: 10 μm. ***B,*** PLA in HeLa cells treated with isoproterenol (1 μM) for 0 (DMSO), 0.5, 3, or 6 hours (*n* = 4 with 71, 52, 48, 56 cells, respectively; Kruskal-Wallis test with Dunn's multiple comparison test). ***C,*** Representative images of TMRM (measuring mitochondrial membrane potential) fluorescent signal in HeLa cells treated with DMSO or CCPA (1 μM each). ***D,*** Representative images of Mito (converted in Image Analyst MKII software) used for length comparison in HeLa cells treated with DMSO or CCPA (1 μM each). Each right panel is magnified images of the corresponding left panel. Scale bars: 10 μm

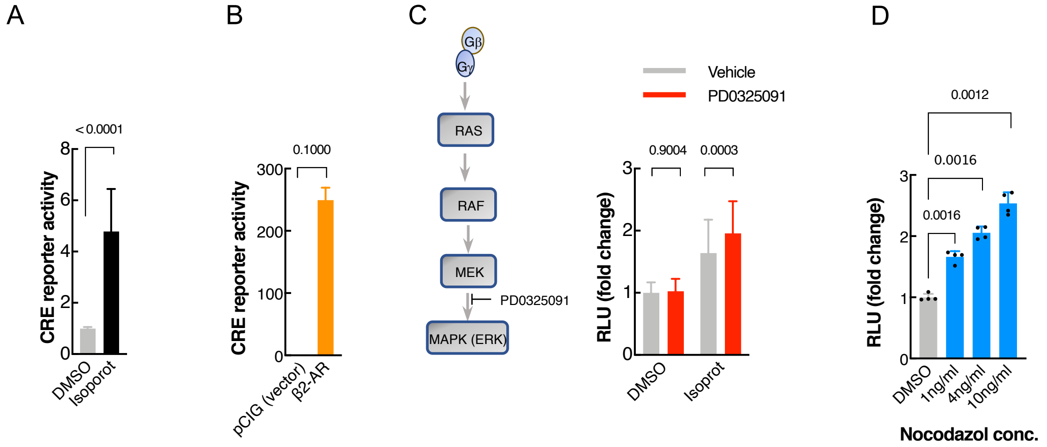


**Figure S3.** ***A,*** cAMP response element (CRE) reporter gene assay in HEK293T cells treated with DMSO or isoproterenol (isoprot, 1 μM). n = 9; Two-tailed unpaired *t*-test. ***B***, cAMP response element (CRE) reporter gene activity in HEK293T cells transfected with pCIG (vector expressing GFP) or β2-AR expression construct. *n* = 3; Mann Whitney test. ***C,*** (Left) Simplified schematic depicting $G_{\beta\chi}$ downstream pathway leading to activation of MAPK. PD0325091 is a chemical inhibitor of MAPK. (Right) Split-Rluc activities (RLU, normalized to the mean of DMSO/Vehicle treated) of the HEK293T cells treated with vehicle or PD0325091 (1 μM, MAPK inhibitor), in the presence of DMSO or isoproterenol (1 μM). *n* = 20. 2-way ANOVA with Sidak's multiple comparisons test. ***D,*** Split-Rluc activities (RLU, normalized to the mean of DMSO) of the HEK293T cells treated with DMSO or indicated concentrations of nocodazole. *n* = 4. 1-way ANOVA with Dunnett's multiple comparisons test. *p* values (if ≤ 0.05, significant) are indicated at the top of each graph. All data: mean ± SD.

**Table S1. Potential hits from the initial screen**

| NCC_STRUC_ID | Drug Name | Target Pathway | Target GPCR | Target AR | Agonist/ Antagonist | sRluc activity |
|---|---|---|---|---|---|---|
| CPD000471620 | Formoterol | GPCR | Adrenergic Rc | beta-2 | agonist | 81826.7 |
| CPD000466295 | Salmeterol | GPCR | Adrenergic Rc | beta-2 | agonist | 81826.7 |
| CPD000058267 | Isoproterenol | GPCR | Adrenergic Rc | beta | agonist | 81826.7 |
| CPD000857209 | Epinephrine | GPCR | Adrenergic Rc | alpha, beta | agonist | 81826.7 |
| CPD000058335 | Triamcinolone (76-25- | Glucocorticoid Rc | | | | 81826.7 |
| CPD000058729 | Duvadilan | GPCR | Adrenergic Rc | beta | agonist | 45628.9 |
| CPD001453712 | Metaproterenol | GPCR | Adrenergic Rc | beta | agonist | 45628.9 |
| CPD000058329 | Fluocinolone | Glucocorticoid Rc | | | | 45628.9 |
| CPD000466308 | Epirubicin | DNA Related | | | | 45628.9 |
| CPD000469221 | Balsalazide | GPCR | Prostaglandin Rc | | | 45628.9 |
| CPD000112594 | Prostaglandin E1 | GPCR | Prostaglandin Rc | | | 45628.9 |
| CPD001496939 | Terbutaline | GPCR | Adrenergic Rc | beta-2 | agonist | 39806.7 |
| CPD000466344 | Topotecan | DNA Related | | | | 39806.7 |
| CPD000466294 | RU 24969 | GPCR | Serotonergic Rc | | | 39806.7 |
| CPD000468732 | CCPA | GPCR | Adenosine Rc | | | 39806.7 |
| CPD000058515 | Artane | GPCR | mAChR | | | 39806.7 |
| CPD001491672 | Phylloquinone | Vitamine K1 | | | | 39806.7 |
| CPD000058513 | Salbutamol | GPCR | Adrenergic Rc | beta-2 | agonist | 33662.2 |
| CPD000058420 | Betaxolol | GPCR | Adrenergic Rc | beta-1 | antagonist | 33662.2 |
| CPD000058383 | Norepinephrine | GPCR | Adrenergic Rc | alpha, beta | agonist | 33662.2 |
| CPD000466920 | Beclomethasone | Glucocorticoid Rc | | | | 33662.2 |
| CPD000058570 | Doxorubicin | DNA Related | | | | 33662.2 |
| CPD000466296 | SB 205607 | GPCR | Opioid Rc | | | 33662.2 |
| CPD000035778 | Hydrochlorothiazide | Transporter | | | | 33662.2 |
| CPD000499581 | Valproic acid | HDAC | | | | 33662.2 |
| CPD000058745 | Clobetasol | Glucocorticoid Rc | | | | 19273.3 |
| CPD001491664 | Amcinonide | Glucocorticoid Rc | | | | 19273.3 |
| CPD000449294 | Capsaicin | Vanilloid Rc | | | | 19273.3 |
| CPD000466355 | Idarubicin | DNA Related | | | | 19273.3 |
| CPD000466319 | Lamivudine | DNA Related | | | | 19273.3 |
| CPD000468734 | PD 81723 | GPCR | Adenosine Rc | | | 19273.3 |
| CPD000469232 | Lofexidine | GPCR | Adrenergic Rc | alpha-2 | agonist | 19273.3 |

**Table S2. GPCR-associated drugs (in NCC library)**

| NCC_Structure_ID | Target GPCR | sRluc activity |
|---|---|---|
| CPD000058832 | Acetylcholine | 14504.4 |
| CPD000058661 | Acetylcholine | 14504.4 |
| CPD000058821 | Acetylcholine | 14084.4 |
| CPD000449327 | Acetylcholine | 13964.4 |
| CPD000469231 | Acetylcholine | 11964.4 |
| CPD000449286 | Acetylcholine | 11773.3 |
| CPD001906769 | Acetylcholine | 11711.1 |
| CPD000058719 | Acetylcholine | 11673.3 |
| CPD000449292 | Acetylcholine | 11620.0 |
| CPD000058660 | Acetylcholine | 11517.8 |
| CPD000471626 | Acetylcholine | 11308.9 |
| CPD000058672 | Acetylcholine | 11186.7 |
| CPD000469196 | Acetylcholine | 11055.6 |
| CPD000058605 | Acetylcholine | 10964.4 |
| CPD000469284 | Acetylcholine | 10846.7 |
| CPD000058817 | Acetylcholine | 10846.7 |
| CPD001906775 | Acetylcholine | 10647.8 |
| CPD000059171 | Acetylcholine | 10613.3 |
| CPD000449267 | Acetylcholine | 10588.9 |
| CPD000059182 | Acetylcholine | 10124.4 |
| CPD000466270 | Acetylcholine | 10033.3 |
| CPD000469282 | Acetylcholine | 5960.0 |
| CPD000057879 | Acetylcholine | 8077.8 |
| CPD000058515 | Acetylcholine | 39806.7 |
| CPD000058523 | Acetylcholine | 13266.7 |
| CPD000471622 | Acetylcholine | 13113.3 |
| CPD000059142 | Acetylcholine | 12557.8 |
| CPD000059053 | Acetylcholine | 12497.8 |
| CPD000058490 | Acetylcholine | 11915.6 |
| CPD000058680 | Acetylcholine | 11593.3 |
| CPD001906768 | Acetylcholine | 11055.6 |
| CPD000058623 | Acetylcholine | 10682.2 |
| CPD000466280 | Acetylcholine | 10601.1 |
| CPD000058440 | Acetylcholine | 10257.8 |
| CPD001496929 | Acetylcholine | 9533.3 |
| CPD000058572 | Acetylcholine | 15577.8 |
| CPD000449277 | Acetylcholine | 12695.6 |
| CPD000471625 | Acetylcholine | 12031.1 |
| CPD000875213 | Acetylcholine | 11786.7 |
| CPD000058736 | Acetylcholine | 12324.4 |
| CPD000468732 | Adenosine | 39806.7 |
| CPD000468734 | Adenosine | 19273.3 |

| | | |
|---|---|---|
| CPD000048468 | Adenosine | 12986.7 |
| CPD000466321 | Adenosine | 11606.7 |
| CPD000466392 | Adenosine | 10846.7 |
| CPD000449316 | Adenosine | 10733.3 |
| CPD000058553 | Adenosine | 10555.6 |
| CPD000449284 | Adenosine | 10148.9 |
| CPD000469631 | Adenosine | 17306.7 |
| CPD000466364 | Adenosine | 15860.0 |
| CPD000469168 | Adenosine | 14800.0 |
| CPD000471620 | Adrenergic | 81826.7 |
| CPD000466295 | Adrenergic | 81826.7 |
| CPD000058267 | Adrenergic | 81826.7 |
| CPD000857209 | Adrenergic | 81826.7 |
| CPD000058729 | Adrenergic | 45628.9 |
| CPD001453712 | Adrenergic | 45628.9 |
| CPD001496939 | Adrenergic | 39806.7 |
| CPD000058513 | Adrenergic | 33662.2 |
| CPD000058420 | Adrenergic | 16197.8 |
| CPD000058383 | Adrenergic | 33662.2 |
| CPD000469232 | Adrenergic | 19273.3 |
| CPD000058800 | Adrenergic | 16197.8 |
| CPD000058463 | Adrenergic | 15577.8 |
| CPD000499584 | Adrenergic | 14606.7 |
| CPD000469190 | Adrenergic | 14555.6 |
| CPD000058525 | Adrenergic | 14412.2 |
| CPD000466362 | Adrenergic | 14320.0 |
| CPD000058423 | Adrenergic | 14320.0 |
| CPD000059167 | Adrenergic | 14320.0 |
| CPD000148117 | Adrenergic | 14146.7 |
| CPD000449302 | Adrenergic | 14146.7 |
| CPD000449280 | Adrenergic | 13964.4 |
| CPD000466276 | Adrenergic | 13964.4 |
| CPD000059120 | Adrenergic | 13964.4 |
| CPD000466346 | Adrenergic | 13822.2 |
| CPD000466347 | Adrenergic | 13660.0 |
| CPD000472527 | Adrenergic | 13528.9 |
| CPD000326828 | Adrenergic | 13222.2 |
| CPD000550486 | Adrenergic | 13222.2 |
| CPD000058388 | Adrenergic | 13148.9 |
| CPD000449301 | Adrenergic | 12813.3 |
| CPD000449266 | Adrenergic | 12733.3 |
| CPD001370753 | Adrenergic | 12695.6 |
| CPD001819784 | Adrenergic | 12466.7 |
| CPD000466340 | Adrenergic | 12324.4 |
| CPD000059081 | Adrenergic | 12310.0 |

| | | |
|---|---|---|
| CPD000058833 | Adrenergic | 12222.2 |
| CPD000058355 | Adrenergic | 12160.0 |
| CPD000449297 | Adrenergic | 12095.6 |
| CPD000036768 | Adrenergic | 12047.8 |
| CPD000058295 | Adrenergic | 12031.1 |
| CPD000449270 | Adrenergic | 12020.0 |
| CPD000059111 | Adrenergic | 11940.0 |
| CPD000149600 | Adrenergic | 11915.6 |
| CPD000058500 | Adrenergic | 11860.0 |
| CPD001906782 | Adrenergic | 11860.0 |
| CPD000058368 | Adrenergic | 11830.0 |
| CPD000238156 | Adrenergic | 11711.1 |
| CPD000469292 | Adrenergic | 11673.3 |
| CPD000058230 | Adrenergic | 11555.6 |
| CPD000469209 | Adrenergic | 11453.3 |
| CPD000036827 | Adrenergic | 11453.3 |
| CPD000449282 | Adrenergic | 11333.3 |
| CPD001453705 | Adrenergic | 11308.9 |
| CPD000097306 | Adrenergic | 11308.9 |
| CPD001496977 | Adrenergic | 11240.0 |
| CPD000059074 | Adrenergic | 11055.6 |
| CPD000469177 | Adrenergic | 10942.2 |
| CPD000059133 | Adrenergic | 10914.4 |
| CPD000471619 | Adrenergic | 10846.7 |
| CPD000058803 | Adrenergic | 10846.7 |
| CPD000466275 | Adrenergic | 10780.0 |
| CPD000058975 | Adrenergic | 10780.0 |
| CPD000058416 | Adrenergic | 10780.0 |
| CPD000059045 | Adrenergic | 10588.9 |
| CPD000449268 | Adrenergic | 10511.1 |
| CPD000058180 | Adrenergic | 10424.4 |
| CPD000449273 | Adrenergic | 10344.4 |
| CPD000469141 | Adrenergic | 10257.8 |
| CPD000058309 | Adrenergic | 10148.9 |
| CPD000058486 | Adrenergic | 10103.3 |
| CPD000469154 | Adrenergic | 10082.2 |
| CPD000058365 | Adrenergic | 10082.2 |
| CPD000058422 | Adrenergic | 9828.9 |
| CPD000058520 | Adrenergic | 9760.0 |
| CPD000466345 | Adrenergic | 9731.1 |
| CPD000326936 | Adrenergic | 9731.1 |
| CPD000471623 | Adrenergic | 9591.1 |
| CPD000058296 | Adrenergic | 9591.1 |
| CPD000469136 | Adrenergic | 9533.3 |
| CPD000058219 | Adrenergic | 9533.3 |

| | | |
|---|---|---|
| CPD000058292 | Adrenergic | 8517.8 |
| CPD000472526 | Adrenergic | 8077.8 |
| CPD000466386 | Angiotensin | 9533.3 |
| CPD000466359 | Angiotensin | 14606.7 |
| CPD000059061 | Angiotensin | 12617.8 |
| CPD000469199 | Angiotensin | 12310.0 |
| CPD000466318 | Angiotensin | 11964.4 |
| CPD001906784 | Angiotensin | 11711.1 |
| CPD000469593 | Angiotensin | 11593.3 |
| CPD000466326 | Angiotensin | 10986.7 |
| CPD000499582 | Angiotensin | 9868.9 |
| CPD000466337 | Angiotensin | 9648.9 |
| CPD000466306 | Angiotensin | 8517.8 |
| CPD000466284 | Cannabinoid | 13028.9 |
| CPD000449274 | Cannabinoid | 12324.4 |
| CPD000466379 | Dopamine | 14555.6 |
| CPD000466274 | Dopamine | 13185.6 |
| CPD000449309 | Dopamine | 12695.6 |
| CPD000449276 | Dopamine | 12617.8 |
| CPD000466292 | Dopamine | 12497.8 |
| CPD000466366 | Dopamine | 12191.1 |
| CPD000058471 | Dopamine | 12160.0 |
| CPD000449283 | Dopamine | 12031.1 |
| CPD000499578 | Dopamine | 11673.3 |
| CPD000449275 | Dopamine | 11646.7 |
| CPD001566944 | Dopamine | 10986.7 |
| CPD000469142 | Dopamine | 10588.9 |
| CPD000058504 | Dopamine | 10588.9 |
| CPD000238142 | Dopamine | 10572.2 |
| CPD000449298 | Dopamine | 10257.8 |
| CPD000058411 | Dopamine | 9984.4 |
| CPD001370746 | Dopamine | 9690.0 |
| CPD000469143 | Dopamine | 14208.9 |
| CPD000058470 | Dopamine | 11308.9 |
| CPD000058957 | Dopamine | 12935.6 |
| CPD000394012 | Dopamine | 13900.0 |
| CPD000466293 | Dopamine | 10942.2 |
| CPD000326935 | Dopamine | 15860.0 |
| CPD000449303 | Dopamine | 12435.6 |
| CPD000058186 | Dopamine | 11517.8 |
| CPD000058380 | Dopamine | 11333.3 |
| CPD000449328 | Dopamine | 10148.9 |
| CPD000466383 | Dopamine | 10192.2 |
| CPD000058465 | Dopamine | 10424.4 |
| CPD000466323 | Dopamine | 13477.8 |

| | | |
|---|---|---|
| CPD000468736 | Dopamine | 11620.0 |
| CPD000058450 | GABA | 13660.0 |
| CPD000550478 | GABA | 13311.1 |
| CPD000449279 | GABA | 12961.1 |
| CPD000149316 | GABA | 12874.4 |
| CPD000058398 | GABA | 12813.3 |
| CPD001491654 | GABA | 12466.7 |
| CPD000058418 | GABA | 12466.7 |
| CPD000238177 | GABA | 12435.6 |
| CPD000238180 | GABA | 12031.1 |
| CPD000469176 | GABA | 12025.6 |
| CPD000469160 | GABA | 11964.4 |
| CPD000596519 | GABA | 11786.7 |
| CPD000058501 | GABA | 11186.7 |
| CPD000466325 | GABA | 10886.7 |
| CPD000059151 | GABA | 10707.8 |
| CPD000058302 | GABA | 10424.4 |
| CPD000010931 | GABA | 10192.2 |
| CPD000469145 | GABA | 9868.9 |
| CPD000469226 | GABA | 6593.3 |
| CPD000469289 | GABA | 6420.0 |
| CPD000499581 | GABA | 33662.2 |
| CPD000058410 | GABA | 18031.1 |
| CPD000466378 | GABA | 15577.8 |
| CPD000058433 | GABA | 14146.7 |
| CPD000058855 | GABA | 10082.2 |
| CPD000449307 | GABA | 11308.9 |
| CPD000059075 | GABA | 10148.9 |
| CPD000058464 | GABA | 10682.2 |
| CPD000857229 | GABA | 18031.1 |
| CPD000471616 | Histamine | 15860.0 |
| CPD001370751 | Histamine | 14800.0 |
| CPD000149358 | Histamine | 13964.4 |
| CPD000058436 | Histamine | 13222.2 |
| CPD000058462 | Histamine | 13131.1 |
| CPD000059100 | Histamine | 12695.6 |
| CPD000466384 | Histamine | 12617.8 |
| CPD000469144 | Histamine | 12222.2 |
| CPD001370748 | Histamine | 12222.2 |
| CPD000469632 | Histamine | 12160.0 |
| CPD001453715 | Histamine | 12095.6 |
| CPD000466315 | Histamine | 11800.0 |
| CPD000058353 | Histamine | 11773.3 |
| CPD000653467 | Histamine | 11453.3 |
| CPD000058255 | Histamine | 10846.7 |

| | | |
|---|---|---|
| CPD000058721 | Histamine | 10846.7 |
| CPD000469183 | Histamine | 10445.6 |
| CPD000058379 | Histamine | 9984.4 |
| CPD000466271 | Histamine | 9788.9 |
| CPD000469220 | Histamine | 9690.0 |
| CPD000471617 | Histamine | 8755.6 |
| CPD000718798 | Histamine | 8517.8 |
| CPD000058961 | Histamine | 7140.0 |
| CPD000469188 | Leukotriene | 14320.0 |
| CPD000466377 | Leukotriene | 13477.8 |
| CPD000466316 | Leukotriene | 13113.3 |
| CPD000469147 | Leukotriene | 12064.4 |
| CPD000058555 | Leukotriene | 10424.4 |
| CPD000059165 | Leukotriene | 9868.9 |
| CPD000466352 | Leukotriene | 9788.9 |
| CPD000449311 | NMDA | 12047.8 |
| CPD000449296 | NMDA | 11915.6 |
| CPD000326694 | NMDA | 12095.6 |
| CPD000058313 | NMDA | 10511.1 |
| CPD000058445 | NMDA | 10445.6 |
| CPD000058999 | NMDA | 10466.7 |
| CPD000466296 | Opioid | 33662.2 |
| CPD000449312 | Opioid | 13861.1 |
| CPD000466389 | Opioid | 13822.2 |
| CPD000058466 | Opioid | 13266.7 |
| CPD000449308 | Opioid | 12222.2 |
| CPD000449320 | Opioid | 11964.4 |
| CPD000058908 | Opioid | 11593.3 |
| CPD000469140 | Opioid | 10942.2 |
| CPD000058766 | Opioid | 10124.4 |
| CPD000449281 | Opioid | 8755.6 |
| CPD000466297 | Opioid | 14800.0 |
| CPD000058382 | Platelets | 10173.3 |
| CPD000466348 | Platelets | 12733.3 |
| CPD000112594 | Prostanoid | 45628.9 |
| CPD000466354 | Prostanoid | 14800.0 |
| CPD000042823 | Prostanoid | 8517.8 |
| CPD000449318 | Prostanoid | 13861.1 |
| CPD000718800 | Prostanoid | 13822.2 |
| CPD000058785 | Prostanoid | 13528.9 |
| CPD000469165 | Prostanoid | 13311.1 |
| CPD000469178 | Prostanoid | 13148.9 |
| CPD000058184 | Prostanoid | 13148.9 |
| CPD000145728 | Prostanoid | 13113.3 |
| CPD000466299 | Prostanoid | 12986.7 |

| | | |
|---|---|---|
| CPD000326718 | Prostanoid | 12813.3 |
| CPD000466327 | Prostanoid | 12733.3 |
| CPD000058991 | Prostanoid | 12047.8 |
| CPD000040181 | Prostanoid | 12020.0 |
| CPD000449291 | Prostanoid | 11860.0 |
| CPD000058443 | Prostanoid | 11800.0 |
| CPD000058746 | Prostanoid | 11692.2 |
| CPD000059146 | Prostanoid | 11240.0 |
| CPD000449290 | Prostanoid | 11186.7 |
| CPD000058206 | Prostanoid | 11186.7 |
| CPD000469285 | Prostanoid | 11108.9 |
| CPD000469164 | Prostanoid | 10942.2 |
| CPD000440694 | Prostanoid | 10733.3 |
| CPD000058188 | Prostanoid | 10588.9 |
| CPD000058461 | Prostanoid | 10344.4 |
| CPD000469594 | Prostanoid | 10211.1 |
| CPD000550473 | Prostanoid | 10082.2 |
| CPD000058835 | Prostanoid | 9648.9 |
| CPD000058286 | Prostanoid | 9271.1 |
| CPD000466331 | Prostanoid | 8517.8 |
| CPD000058723 | Prostanoid | 6593.3 |
| CPD000058715 | Prostanoid | 6420.0 |
| CPD000466294 | Serotonin | 39806.7 |
| CPD000059115 | Serotonin | 5960.0 |
| CPD000466283 | Serotonin | 18031.1 |
| CPD000469200 | Serotonin | 15860.0 |
| CPD000466268 | Serotonin | 13477.8 |
| CPD000449310 | Serotonin | 13477.8 |
| CPD000058254 | Serotonin | 13477.8 |
| CPD001307702 | Serotonin | 13113.3 |
| CPD000469228 | Serotonin | 12961.1 |
| CPD000525252 | Serotonin | 12935.6 |
| CPD000012114 | Serotonin | 12874.4 |
| CPD000469138 | Serotonin | 12714.4 |
| CPD000059131 | Serotonin | 12714.4 |
| CPD000469203 | Serotonin | 12295.6 |
| CPD000469191 | Serotonin | 12160.0 |
| CPD000058507 | Serotonin | 12064.4 |
| CPD000469211 | Serotonin | 11964.4 |
| CPD000112269 | Serotonin | 11860.0 |
| CPD000449305 | Serotonin | 11800.0 |
| CPD000466269 | Serotonin | 11646.7 |
| CPD000469179 | Serotonin | 11453.3 |
| CPD000469156 | Serotonin | 11186.7 |
| CPD000466298 | Serotonin | 11108.9 |

| | | |
|---|---|---|
| CPD000471618 | Serotonin | 11108.9 |
| CPD001227191 | Serotonin | 11021.1 |
| CPD000465669 | Serotonin | 10866.7 |
| CPD000469233 | Serotonin | 10613.3 |
| CPD000449271 | Serotonin | 10211.1 |
| CPD000466277 | Serotonin | 10211.1 |
| CPD000469158 | Serotonin | 10173.3 |
| CPD000058431 | Serotonin | 9788.9 |
| CPD000466272 | Serotonin | 9648.9 |
| CPD000058452 | Serotonin | 7608.9 |
| CPD000059105 | Serotonin | 6593.3 |
| CPD000449269 | Serotonin | 4731.1 |
| CPD000449272 | Serotonin | 12222.2 |
| CPD000449287 | Serotonin | 11055.6 |
| CPD000058269 | Serotonin | 6420.0 |
| CPD001453706 | Serotonin | 10707.8 |
| CPD000469181 | Serotonin | 10613.3 |
| CPD000058926 | Serotonin | 12733.3 |
| CPD000449299 | Serotonin | 13028.9 |

**Table S3. AR-associated drugs (in NCC library)**

| NCC_STRUC_ID | Target AR | Agonist/ Antagonist | sRluc activity |
|---|---|---|---|
| CPD000469232 | alpha | agonist | 19273.3 |
| CPD000058833 | alpha | agonist | 12222.2 |
| CPD000059111 | alpha | agonist | 11940.0 |
| CPD000058219 | alpha | agonist | 9533.3 |
| CPD000058292 | alpha | agonist | 8517.8 |
| CPD000499584 | alpha | agonist | 14606.7 |
| CPD000466276 | alpha | agonist | 13964.4 |
| CPD001370753 | alpha | agonist | 12695.6 |
| CPD000058355 | alpha | agonist | 12160.0 |
| CPD000469209 | alpha | agonist | 11453.3 |
| CPD000058803 | alpha | agonist | 10846.7 |
| CPD001906782 | alpha | antagonist | 11860.0 |
| CPD000059133 | alpha | antagonist | 10914.4 |
| CPD000059045 | alpha | antagonist | 10588.9 |
| CPD000058180 | alpha | antagonist | 10424.4 |
| CPD000058422 | alpha | antagonist | 9828.9 |
| CPD000058520 | alpha | antagonist | 9760.0 |
| CPD000058525 | alpha | antagonist | 14412.2 |
| CPD000466362 | alpha | antagonist | 14320.0 |
| CPD000466346 | alpha | antagonist | 13822.2 |
| CPD000449301 | alpha | antagonist | 12813.3 |
| CPD000466340 | alpha | antagonist | 12324.4 |
| CPD000097306 | alpha | antagonist | 11308.9 |
| CPD000466275 | alpha | antagonist | 10780.0 |
| CPD000449268 | alpha | antagonist | 10511.1 |
| CPD000058309 | alpha | antagonist | 10148.9 |
| CPD001819784 | alpha | antagonist | 12466.7 |
| CPD000471623 | alpha | antagonist | 9591.1 |
| CPD000469190 | alpha | antagonist | 14555.6 |
| CPD000449302 | alpha | antagonist | 14146.7 |
| CPD000466347 | alpha | antagonist | 13660.0 |
| CPD000550486 | alpha | antagonist | 13222.2 |
| CPD000149600 | alpha | antagonist | 11915.6 |
| CPD000058365 | alpha | antagonist | 10082.2 |
| CPD000857209 | alpha, beta | agonist | 81826.7 |
| CPD000058383 | alpha, beta | agonist | 33662.2 |
| CPD000059081 | alpha, beta | agonist | 12310.0 |
| CPD000058463 | alpha, beta | antagonist | 15577.8 |
| CPD000466345 | alpha, beta | antagonist | 9731.1 |
| CPD000449280 | alpha, beta | antagonist | 13964.4 |
| CPD000058296 | alpha, beta | antagonist | 9591.1 |

| | | | |
|---|---|---|---|
| CPD000058267 | beta | agonist | 81826.7 |
| CPD000058729 | beta | agonist | 45628.9 |
| CPD001453712 | beta | agonist | 45628.9 |
| CPD000471620 | beta | agonist | 81826.7 |
| CPD000466295 | beta | agonist | 81826.7 |
| CPD001496939 | beta | agonist | 39806.7 |
| CPD000058513 | beta | agonist | 33662.2 |
| CPD000058420 | beta | antagonist | 33662.2 |
| CPD000058800 | beta | antagonist | 16197.8 |
| CPD000059167 | beta | antagonist | 14320.0 |
| CPD000059120 | beta | antagonist | 13964.4 |
| CPD000326828 | beta | antagonist | 13222.2 |
| CPD000058388 | beta | antagonist | 13148.9 |
| CPD000036768 | beta | antagonist | 12047.8 |
| CPD000471619 | beta | antagonist | 10846.7 |
| CPD000058975 | beta | antagonist | 10780.0 |
| CPD000469141 | beta | antagonist | 10257.8 |
| CPD001453705 | beta | antagonsit | 11308.9 |

**Table S4. mRNA seq data for adrenergic receptors**

| | TPM (transcripts per million) | | | TPM (transcripts per million) | |
|---|---|---|---|---|---|
| **Gene name** | **HEK293T** | **HeLa** | **Gene name** | **C2C12** | **Neuro2A** |
| **ACKR1** | NA | NA | **Ackr1** | NA | 0.411135 |
| **ACKR2** | 0.0671921 | 0.00611861 | **Ackr2** | 0.0290849 | 0.0274002 |
| **ACKR3** | NA | 10.7309 | **Ackr3** | 10.5631 | 0.0218309 |
| **ACKR4** | 2.40158 | 1.43155 | **Ackr4** | 4.28392 | 0.273759 |
| **ADCYAP1R1** | 0.412282 | 0.0204383 | **Adcyap1r1** | NA | 10.1754 |
| **ADGRA1** | 0.014162 | 0.00209562 | **Adgra1** | 0.0280815 | 0.349205 |
| **ADGRA2** | 1.06185 | 1.11038 | **Adgra2** | 17.5953 | 3.67736 |
| **ADGRA3** | 1.52622 | 1.54665 | **Adgra3** | 4.41482 | 1.44618 |
| **ADGRB1** | 0.0186445 | 0.276368 | **Adgrb1** | NA | 0.139848 |
| **ADGRB2** | 0.99388 | 2.11289 | **Adgrb2** | 0.205729 | 0.369166 |
| **ADGRB3** | 0.55272 | 0.000928321 | **Adgrb3** | NA | 0.0175584 |
| **ADGRD1** | 0.00229335 | 0.0678718 | **Adgrd1** | 0.205819 | 4.37884 |
| **ADGRD2** | 0.00678628 | 0.569381 | **Adgre1** | 0.0296354 | 0.0217767 |
| **ADGRE1** | 0.00116175 | 0.00722024 | **Adgre5** | 20.6852 | 0.871518 |
| **ADGRE2** | 0.0053239 | 6.33217 | **Adgrf1** | NA | 0.00896804 |
| **ADGRE3** | 1.52864 | 9.58569 | **Adgrf2** | 0.0428473 | 0.292145 |
| **ADGRE4P** | 0.00135432 | NA | **Adgrf3** | 0.170553 | 0.0226834 |
| **ADGRE5** | 0.333102 | 16.6925 | **Adgrf4** | NA | 0.189065 |
| **ADGRF1** | NA | 0.00570313 | **Adgrf5** | 0.0398179 | 0.0858076 |
| **ADGRF2** | 0.00742829 | NA | **Adgrg1** | 4.8713 | 0.190748 |
| **ADGRF3** | 0.386397 | 0.331062 | **Adgrg2** | 0.119437 | 0.0467386 |
| **ADGRF4** | 0.00509844 | 0.0475298 | **Adgrg3** | 1.90625 | 0.134055 |
| **ADGRF5** | 0.00119993 | 0.0031073 | **Adgrg4** | 0.00656975 | 0.00530503 |
| **ADGRG1** | 0.00111871 | 4.58533 | **Adgrg5** | 0.214064 | 0.0142352 |
| **ADGRG2** | 0.161309 | 0.208143 | **Adgrg6** | 4.31215 | 3.71776 |
| **ADGRG3** | NA | 0.241309 | **Adgrg7** | NA | 0.0166323 |
| **ADGRG4** | 0.00318041 | 0.00164717 | **Adgrl1** | 11.9194 | 17.6022 |
| **ADGRG5** | 0.00125308 | 0.0986458 | **Adgrl2** | 5.19926 | 1.72021 |
| **ADGRG6** | 0.37275 | 0.133109 | **Adgrl3** | 0.298114 | 0.722868 |
| **ADGRG7** | 0.00572431 | 0.00370586 | **Adgrl4** | 0.0264586 | 0.0123546 |
| **ADGRL1** | NA | NA | **Adgrv1** | 0.00687546 | 0.0687879 |
| **ADGRL1** | 1.9325 | 1.72728 | **Adora1** | 14.243 | 0.173259 |
| **ADGRL2** | 0.00877483 | 0.00565755 | **Adora2a** | 0.406292 | 8.11576 |
| **ADGRL3** | 0.61066 | 0.000653147 | **Adora2b** | 5.09656 | NA |
| **ADGRL4** | 0.00125292 | 0.0629437 | **Adora3** | NA | 0.0292967 |
| **ADGRV1** | 0.776475 | 0.0175391 | **Adra1a** | 0.00181462 | 0.00923136 |
| **ADORA1** | 0.0640502 | 0.0514172 | **Adra1b** | 0.0915609 | 0.010461 |
| **ADORA2A** | 0.602473 | 0.759269 | **Adra1d** | 2.88152 | 0.0508989 |
| **ADORA2B** | 1.27165 | 1.93039 | **Adra2a** | NA | 0.31737 |
| **ADORA3** | NA | NA | **Adra2b** | NA | 1.47609 |
| **ADRA1A** | 0.0128956 | NA | **Adra2c** | NA | 0.0809359 |
| **ADRA1B** | 0.0142333 | 0.159227 | **Adrb1** | 0.0677892 | 0.0447038 |
| **ADRA1D** | 0.00643274 | 0.0666319 | **Adrb2** | 33.8209 | NA |
| **ADRA2A** | 0.0633666 | NA | **Adrb3** | 0.173753 | 0.0140304 |
| **ADRA2B** | 0.0498781 | 0.137773 | **Agtr2** | NA | 0.0301692 |
| **ADRA2C** | 5.97859 | 5.17038 | **Aplnr** | 0.666774 | 0.0565337 |
| **ADRB1** | 0.465069 | 1.36141 | **Avpr1a** | NA | 0.0281413 |
| **ADRB2** | 53.3648 | 2.82853 | **Avpr1b** | NA | 0.114702 |

| | | | | | |
|---|---|---|---|---|---|
| **ADRB3** | NA | NA | **Avpr2** | 0.217559 | 0.471402 |
| **AGTR1** | 0.134517 | 0.00985204 | **Bdkrb1** | 4.73498 | 0.118952 |
| **AGTR2** | NA | 0.030088 | **Bdkrb2** | 1.98779 | 2.53243 |
| **APLNR** | NA | 0.191354 | **Brs3** | NA | NA |
| **AVPR1A** | 0.0800601 | NA | **C3ar1** | 0.478302 | 0.015449 |
| **AVPR1B** | 0.0228353 | NA | **C5ar1** | 0.0481794 | 0.0980394 |
| **AVPR2** | 0.0397647 | 2.64984 | **C5ar2** | 0.0128861 | NA |
| **BDKRB1** | NA | 0.0798259 | **Calcr** | 0.00468096 | 0.0582096 |
| **BDKRB2** | 0.197633 | 0.538379 | **Calcrl** | 1.56514 | 0.0956025 |
| **BRS3** | 0.147463 | NA | **Casr** | NA | 0.00611045 |
| **C3AR1** | 0.0193726 | 0.0401332 | **Cckar** | NA | 0.0140395 |
| **C5AR1** | 0.0441064 | 0.572074 | **Cckbr** | 0.0409537 | 0.0077163 |
| **C5AR2** | 0.0121643 | 5.84222 | **Ccr1** | 0.037546 | NA |
| **CALCR** | 0.00245398 | NA | **Ccr10** | 1.76387 | 3.61776 |
| **CALCRL** | 0.0761653 | 0.00119536 | **Ccr2** | NA | 0.042035 |
| **CASR** | NA | 0.00235828 | **Ccr3** | NA | NA |
| **CCKAR** | NA | NA | **Ccr4** | 0.00844599 | 0.17664 |
| **CCKBR** | 0.074805 | NA | **Ccr5** | 0.047146 | 0.0239842 |
| **CCR1** | NA | NA | **Ccr6** | 0.0116896 | 0.0991128 |
| **CCR10** | 0.53764 | 5.2779 | **Ccr7** | 0.0679948 | 0.134518 |
| **CCR2** | NA | NA | **Ccr8** | 0.0883064 | NA |
| **CCR3** | 0.00178801 | 0.0265463 | **Ccr9** | 1.58616 | 12.525 |
| **CCR4** | NA | 0.108204 | **Ccrl2** | 1.5039 | 0.252724 |
| **CCR5** | NA | NA | **Celsr1** | 0.0118511 | 0.024734 |
| **CCR6** | 0.0881316 | 0.02054 | **Celsr2** | NA | 1.93016 |
| **CCR7** | NA | 1.23996 | **Celsr3** | 0.101803 | 12.6988 |
| **CCR8** | 0.061883 | 0.0320499 | **Chrm1** | 0.0373716 | 0.225324 |
| **CCR9** | 0.0663445 | 0.0152714 | **Chrm2** | 0.000878779 | 0.00596071 |
| **CCRL2** | NA | 0.087313 | **Chrm3** | 0.00177855 | 0.196755 |
| **CELSR1** | 0.499838 | 2.56748 | **Chrm4** | 1.88405 | 15.5386 |
| **CELSR2** | 1.65504 | 14.6301 | **Chrm5** | 0.154246 | 0.0435935 |
| **CELSR3** | 2.55111 | 17.0353 | **Cmklr1** | 0.715827 | 0.0549255 |
| **CHRM1** | NA | 0.0145444 | **Cnr1** | 4.17697 | 17.3266 |
| **CHRM2** | 0.100533 | NA | **Cnr2** | NA | 0.039328 |
| **CHRM3** | 0.0216107 | 0.0973624 | **Crhr1** | NA | 0.486826 |
| **CHRM4** | 0.447351 | 1.64288 | **Crhr2** | 0.0114802 | 0.00162228 |
| **CHRM5** | 0.195597 | 0.0611029 | **Cx3cr1** | 6.74777 | 1.03732 |
| **CMKLR1** | NA | NA | **Cxcr1** | NA | NA |
| **CNR1** | 0.201771 | NA | **Cxcr2** | NA | 0.019208 |
| **CNR2** | 0.0100389 | 0.0118841 | **Cxcr3** | NA | NA |
| **CRHR1** | NA | NA | **Cxcr4** | 9.96798 | NA |
| **CRHR1** | NA | NA | **Cxcr5** | 0.288044 | 0.20178 |
| **CRHR1** | 0.0882128 | NA | **Cxcr6** | 0.839863 | 0.0923084 |
| **CRHR2** | 0.0395315 | 0.184925 | **Cysltr1** | NA | NA |
| **CX3CR1** | 0.0505316 | NA | **Cysltr2** | 0.0576947 | 0.275717 |
| **CXCR1** | NA | NA | **Drd1** | NA | 0.123292 |
| **CXCR2** | NA | 0.0425654 | **Drd2** | 0.00182558 | 0.00515951 |
| **CXCR3** | 0.0710128 | NA | **Drd3** | 0.00715863 | 0.0890204 |
| **CXCR4** | 4.20335 | 19.1741 | **Drd4** | 0.110626 | 0.140694 |
| **CXCR5** | 0.106131 | 23.1592 | **Drd5** | NA | 0.0442297 |
| **CXCR6** | 1.78908 | 0.154431 | **Ednra** | 0.843089 | 0.00794254 |
| **CYSLTR1** | 0.00218589 | NA | **Ednrb** | NA | 0.0468171 |

| | | | | | |
|---|---|---|---|---|---|
| **CYSLTR2** | 0.00213679 | 0.00885335 | **F2r** | 70.0968 | 41.9615 |
| **DRD1** | NA | NA | **F2rl1** | NA | 0.00516494 |
| **DRD2** | 0.0102281 | NA | **F2rl2** | 0.418957 | 17.5242 |
| **DRD3** | 0.00171103 | NA | **F2rl3** | 1.35992 | 0.663867 |
| **DRD4** | NA | NA | **Ffar1** | NA | NA |
| **DRD4** | 0.107242 | 1.38855 | **Ffar2** | NA | NA |
| **DRD5** | NA | NA | **Ffar3** | NA | 0.0376993 |
| **EDNRA** | 0.531609 | 1.98593 | **Ffar4** | NA | 0.158119 |
| **EDNRB** | 0.122824 | 0.00397576 | **Fpr1** | 0.0990521 | NA |
| **F2R** | 3.71295 | 5.43684 | **Fpr2** | 0.120724 | 0.0113731 |
| **F2RL1** | 3.50702 | 2.1796 | **Fpr3** | 0.0176537 | 0.353411 |
| **F2RL2** | 0.240159 | NA | **Fshr** | 0.00114331 | 0.0739958 |
| **F2RL3** | NA | 0.0705667 | **Fzd1** | 105.367 | 0.398605 |
| **FFAR1** | NA | NA | **Fzd10** | 0.0758892 | 0.021448 |
| **FFAR2** | NA | NA | **Fzd2** | 57.4348 | 0.837309 |
| **FFAR3** | NA | NA | **Fzd3** | 0.533685 | 2.26795 |
| **FFAR4** | NA | 0.351785 | **Fzd4** | 11.0258 | 4.82702 |
| **FPR1** | 0.0041704 | 0.0043198 | **Fzd5** | 39.3239 | 6.21095 |
| **FPR2** | 0.0132821 | 0.013758 | **Fzd6** | 4.64439 | 1.4546 |
| **FPR3** | 0.0396016 | NA | **Fzd7** | 53.3606 | 32.5613 |
| **FSHR** | 0.00223604 | 0.000992632 | **Fzd8** | 4.64462 | 0.0700094 |
| **FZD1** | 3.98434 | 2.89911 | **Fzd9** | 2.15124 | 0.425593 |
| **FZD10** | 1.42297 | 32.7759 | **Gabbr1** | 20.3612 | 3.95453 |
| **FZD2** | 4.03269 | 23.5639 | **Gabbr2** | NA | 0.0188241 |
| **FZD3** | 3.16524 | 0.552535 | **Galr1** | NA | NA |
| **FZD4** | 7.82272 | 12.8324 | **Galr2** | 0.698813 | 0.929413 |
| **FZD5** | 8.55875 | 5.40804 | **Galr3** | 0.221003 | 0.208202 |
| **FZD6** | 6.57078 | 5.44383 | **Gcgr** | 0.208311 | 0.0336419 |
| **FZD7** | 13.9492 | 7.43889 | **Ghrhr** | NA | NA |
| **FZD8** | 5.23009 | 6.82315 | **Ghsr** | 0.0185165 | NA |
| **FZD9** | 1.96702 | 6.68297 | **Gipr** | 0.489997 | 2.07726 |
| **GABBR1** | NA | NA | **Glp1r** | 0.563206 | 2.35901 |
| **GABBR1** | NA | NA | **Glp2r** | 0.328397 | 0.0902697 |
| **GABBR1** | NA | NA | **Gna12** | 5.04073 | 2.53181 |
| **GABBR1** | NA | NA | **Gna13** | 15.065 | 18.3693 |
| **GABBR1** | NA | NA | **Gna14** | 0.00845337 | 0.0314567 |
| **GABBR1** | NA | NA | **Gna15** | 0.180029 | 0.213061 |
| **GABBR1** | 0.47765 | 0.780732 | **Gnai1** | 0.0362626 | 0.0541713 |
| **GABBR2** | 0.0559262 | 0.0116465 | **Gnai2** | 147.998 | 12.1378 |
| **GALR1** | NA | NA | **Gnal** | 1.14441 | 0.293256 |
| **GALR2** | 0.0891546 | 0.138523 | **Gnao1** | 7.7029 | 3.62646 |
| **GALR3** | 0.0876185 | 0.0907574 | **Gnaq** | 1.65148 | 5.21112 |
| **GCGR** | NA | NA | **Gnas** | 88.98 | 48.5078 |
| **GCGR** | 0.0062215 | 4.46596 | **Gnat1** | 0.0470776 | 0.0532208 |
| **GHRHR** | 0.00450297 | NA | **Gnat2** | 1.75478 | 2.12892 |
| **GHSR** | 0.0356851 | NA | **Gnat3** | 0.00645812 | 0.0304202 |
| **GIPR** | 0.253695 | 5.20641 | **Gnaz** | 0.225656 | 1.45212 |
| **GLP1R** | 0.00289019 | NA | **Gnrhr** | NA | NA |
| **GLP2R** | NA | 7.13428 | **Gpbar1** | NA | 1.88044 |
| **GNA12** | 0.87518 | 3.39553 | **Gper1** | 11.4568 | 0.749527 |
| **GNA13** | 7.74662 | 7.70489 | **Gpr1** | 0.0348438 | 0.048144 |
| **GNA14** | 0.0103669 | 0.00989057 | **Gpr101** | NA | 0.0294948 |

| | | | | | |
|---|---|---|---|---|---|
| **GNA15** | 0.00443409 | 0.296244 | **Gpr107** | 7.54281 | 2.90739 |
| **GNAI1** | 0.37166 | 0.530017 | **Gpr119** | 0.0379446 | NA |
| **GNAI2** | 4.25137 | 30.8026 | **Gpr12** | NA | 3.55643 |
| **GNAL** | 1.0252 | 1.548 | **Gpr132** | 0.0284213 | NA |
| **GNAO1** | 0.0529181 | 1.52942 | **Gpr135** | 0.782983 | 1.6818 |
| **GNAQ** | 1.19176 | 1.04051 | **Gpr137** | 22.4674 | 15.4727 |
| **GNAS** | 20.9845 | 45.9669 | **Gpr139** | 0.0112475 | 0.00317879 |
| **GNAT1** | NA | 0.282336 | **Gpr141** | 0.0992168 | 0.0177593 |
| **GNAT2** | 2.02671 | 1.6222 | **Gpr142** | 0.268018 | 0.0534692 |
| **GNAT3** | 0.0046004 | NA | **Gpr143** | 0.00922846 | 0.0234735 |
| **GNAZ** | 2.29504 | 0.754277 | **Gpr146** | 5.65246 | 1.54911 |
| **GNRHR** | 1.05672 | 0.784998 | **Gpr149** | 0.194552 | 0.00549848 |
| **GPBAR1** | 0.253343 | 0.0728942 | **Gpr15** | 0.301147 | NA |
| **GPER1** | 0.3388 | 7.21614 | **Gpr150** | NA | 0.0324818 |
| **GPR1** | NA | NA | **Gpr151** | 0.347968 | 0.118012 |
| **GPR1** | 0.0388267 | 0.210026 | **Gpr152** | 0.0414242 | 0.0117074 |
| **GPR101** | NA | NA | **Gpr153** | 5.56305 | 2.2093 |
| **GPR107** | 3.32241 | 8.076 | **Gpr156** | 0.0731507 | 0.0589593 |
| **GPR119** | NA | NA | **Gpr157** | 1.6874 | 0.0150204 |
| **GPR12** | 0.0329961 | NA | **Gpr158** | 0.00319615 | 0.051639 |
| **GPR132** | 0.00765682 | 0.21414 | **Gpr160** | 0.233226 | 0.0219717 |
| **GPR135** | 0.908525 | 2.66024 | **Gpr161** | 2.98769 | 0.434579 |
| **GPR137** | 3.14219 | 11.5587 | **Gpr162** | 3.27221 | 5.07067 |
| **GPR139** | NA | NA | **Gpr17** | 0.705746 | 0.440911 |
| **GPR141** | 0.00245811 | 0.0169744 | **Gpr171** | NA | NA |
| **GPR142** | NA | NA | **Gpr173** | 4.22904 | 2.40988 |
| **GPR143** | 0.006049 | 0.110694 | **Gpr174** | 0.012063 | 0.018751 |
| **GPR146** | 0.492734 | 3.53377 | **Gpr176** | 2.09097 | 0.0709292 |
| **GPR148** | NA | NA | **Gpr179** | 0.0494045 | 0.230387 |
| **GPR149** | 0.0135483 | 0.00534614 | **Gpr18** | 0.190603 | 0.0769552 |
| **GPR15** | NA | NA | **Gpr182** | 1.74314 | 0.563028 |
| **GPR150** | NA | NA | **Gpr183** | 0.0195049 | 0.0551254 |
| **GPR151** | 0.338292 | NA | **Gpr19** | 1.89041 | 2.93141 |
| **GPR152** | NA | NA | **Gpr20** | 0.0382328 | 0.0756382 |
| **GPR153** | 1.80603 | 9.25179 | **Gpr21** | 0.461872 | 0.163169 |
| **GPR156** | 0.0862185 | 0.951201 | **Gpr22** | NA | 5.34534 |
| **GPR157** | 1.21628 | 3.13415 | **Gpr25** | NA | NA |
| **GPR158** | 0.00460052 | 0.0735649 | **Gpr26** | NA | 0.142176 |
| **GPR160** | 1.99396 | 1.1079 | **Gpr27** | NA | 0.233521 |
| **GPR161** | 2.0774 | 0.900151 | **Gpr3** | 0.115721 | 5.29827 |
| **GPR162** | 0.802255 | 0.404062 | **Gpr33** | NA | NA |
| **GPR17** | 0.0816187 | 4.48076 | **Gpr34** | 0.140616 | 0.0953788 |
| **GPR171** | 0.981371 | NA | **Gpr35** | 0.32976 | 0.0706341 |
| **GPR173** | 0.00386149 | 0.553976 | **Gpr37** | 0.029773 | 0.0476824 |
| **GPR174** | NA | NA | **Gpr37l1** | 0.0331149 | 0.393079 |
| **GPR176** | 0.74089 | 1.01152 | **Gpr39** | 0.329491 | 0.0123926 |
| **GPR179** | NA | NA | **Gpr4** | 0.264931 | 0.155741 |
| **GPR179** | 0.0760753 | 0.0656672 | **Gpr45** | 0.0029865 | 0.098754 |
| **GPR18** | 1.2061 | 0.162953 | **Gpr50** | NA | 0.0124675 |
| **GPR182** | 0.233 | 0.273526 | **Gpr52** | 0.602663 | 0.0425816 |
| **GPR183** | 1.00816 | 0.445425 | **Gpr55** | NA | 0.00921631 |
| **GPR19** | 0.186169 | 0.151387 | **Gpr6** | NA | 0.120755 |

| | | | | | |
|---|---|---|---|---|---|
| **GPR20** | NA | NA | **Gpr61** | 0.18793 | 0.0743587 |
| **GPR20** | 0.0454468 | 0.841464 | **Gpr62** | 6.8615 | 2.18603 |
| **GPR21** | 0.778702 | 0.0467594 | **Gpr63** | 0.820654 | 0.00651047 |
| **GPR22** | NA | NA | **Gpr65** | 0.0307915 | 0.0261071 |
| **GPR22** | 1.16025 | 0.314493 | **Gpr68** | 1.43209 | 1.90206 |
| **GPR25** | NA | NA | **Gpr75** | 0.206221 | 0.566174 |
| **GPR26** | NA | NA | **Gpr82** | 0.243755 | 0.0688908 |
| **GPR27** | 15.4206 | NA | **Gpr83** | NA | 0.0131769 |
| **GPR3** | 0.327168 | 7.63495 | **Gpr84** | 0.103717 | 0.0586257 |
| **GPR31** | NA | NA | **Gpr85** | 0.764426 | 7.23112 |
| **GPR32** | 0.0484238 | NA | **Gpr87** | NA | NA |
| **GPR33** | NA | NA | **Gpr88** | 0.320311 | 0.0362109 |
| **GPR34** | 0.355244 | 0.298976 | **Gprc5a** | 6.95571 | 0.168239 |
| **GPR35** | 0.123708 | 0.211924 | **Gprc5b** | 27.9859 | 0.213647 |
| **GPR37** | 0.224392 | 13.1323 | **Gprc5c** | 2.46456 | 0.0389673 |
| **GPR37L1** | 0.207212 | 0.667753 | **Gprc5d** | 0.381205 | 0.0510334 |
| **GPR39** | 0.308081 | 0.137952 | **Gprc6a** | NA | 0.00837863 |
| **GPR4** | 0.00987224 | 0.132937 | **Grm1** | 0.0230273 | 0.141946 |
| **GPR42** | NA | NA | **Grm2** | NA | 0.0751374 |
| **GPR45** | 0.142492 | 0.368992 | **Grm3** | 0.00205851 | 0.0113447 |
| **GPR50** | 0.151044 | 0.0260759 | **Grm4** | 0.00249088 | 0.00915174 |
| **GPR52** | 1.20414 | 0.0804693 | **Grm5** | 0.00470225 | 0.0322748 |
| **GPR55** | NA | 0.117239 | **Grm6** | NA | NA |
| **GPR6** | NA | NA | **Grm7** | 0.023148 | 0.010059 |
| **GPR61** | 0.84237 | 0.372883 | **Grm8** | 0.00157724 | 0.0286909 |
| **GPR62** | 0.0579989 | 0.36046 | **Grpr** | 0.0999686 | 0.00584553 |
| **GPR63** | 2.46491 | 0.0205549 | **Hcar1** | NA | 0.0848779 |
| **GPR65** | NA | NA | **Hcar2** | 0.127793 | 0.0722342 |
| **GPR68** | 0.0143615 | 0.119008 | **Hcrtr1** | 0.0269758 | 0.0609918 |
| **GPR75** | 1.41955 | 3.72562 | **Hcrtr2** | 0.00251732 | 0.00853743 |
| **GPR78** | 0.387604 | 0.5235 | **Hrh1** | 0.0606768 | 0.200884 |
| **GPR82** | 0.330486 | 0.117674 | **Hrh2** | NA | 0.00949414 |
| **GPR83** | 0.132567 | 0.0580952 | **Hrh3** | NA | 1.94244 |
| **GPR84** | NA | 0.591959 | **Hrh4** | NA | 0.0350877 |
| **GPR85** | 0.598368 | 0.026948 | **Htr1a** | NA | 0.139909 |
| **GPR87** | 0.194607 | 0.0167982 | **Htr1b** | 1.91504 | 0.0251737 |
| **GPR88** | 0.0791267 | NA | **Htr1d** | 0.0590754 | 0.016696 |
| **GPRC5A** | 0.519478 | 179.4 | **Htr1f** | 0.0142979 | 0.0392545 |
| **GPRC5B** | 1.24038 | 0.557113 | **Htr2a** | 0.0808302 | 0.0111684 |
| **GPRC5C** | 0.414969 | 2.81886 | **Htr2b** | 1.00028 | 0.425398 |
| **GPRC5D** | 0.0324188 | 10.9975 | **Htr2c** | 0.135525 | 0.00682912 |
| **GPRC6A** | 0.0149476 | NA | **Htr4** | 0.0100159 | 0.0105141 |
| **GRM1** | 0.00824415 | 0.0080737 | **Htr5a** | NA | 0.0161606 |
| **GRM2** | 0.0691882 | 0.00551283 | **Htr6** | 0.0255797 | 0.583173 |
| **GRM3** | 0.108177 | 0.0483928 | **Htr7** | 0.0112366 | 0.0162314 |
| **GRM4** | 0.000897208 | 0.136614 | **Kiss1r** | 3.09288 | 1.57736 |
| **GRM5** | 0.00186089 | 0.000680313 | **Lgr4** | 3.70045 | 0.466818 |
| **GRM6** | NA | NA | **Lgr5** | 0.00897068 | 0.785441 |
| **GRM7** | 0.00120176 | 0.00746885 | **Lgr6** | 0.181983 | 0.00514326 |
| **GRM8** | 0.00588472 | 0.00265703 | **Lhcgr** | NA | 0.00830822 |
| **GRPR** | 0.00820589 | 0.495117 | **Lpar1** | 3.06418 | 0.0117504 |
| **HCAR1** | 0.051541 | NA | **Lpar2** | 1.3222 | 4.17079 |

| | | | | | |
|---|---|---|---|---|---|
| **HCAR2** | NA | 7.42854 | **Lpar3** | 0.308186 | 0.00106872 |
| **HCAR3** | 0.119552 | 6.25367 | **Lpar4** | 15.0574 | 0.0367537 |
| **HCRTR1** | 13.6587 | 7.28215 | **Lpar5** | NA | 0.349656 |
| **HCRTR2** | 0.00104648 | NA | **Lpar6** | 5.77777 | 3.1755 |
| **HRH1** | 0.00680265 | 1.98657 | **Ltb4r2** | NA | 0.0751345 |
| **HRH2** | 0.026805 | 0.00120718 | **Mas1** | 0.337176 | 0.0669631 |
| **HRH3** | NA | 0.203435 | **Mc1r** | 0.0677954 | 0.36405 |
| **HRH4** | 0.0540567 | NA | **Mc2r** | 0.187065 | 1.68558 |
| **HTR1A** | NA | NA | **Mc3r** | NA | NA |
| **HTR1B** | NA | NA | **Mc4r** | 2.0678 | 0.0248684 |
| **HTR1D** | 0.476859 | 8.06024 | **Mc5r** | 0.203372 | 0.0287388 |
| **HTR1E** | NA | 0.000804164 | **Mchr1** | 1.31646 | 0.160891 |
| **HTR1F** | 0.09785 | NA | **Mrgprd** | NA | NA |
| **HTR2A** | 0.0103156 | NA | **Mrgpre** | 11.7735 | 1.28328 |
| **HTR2B** | 1.07454 | 0.162791 | **Mrgprf** | 14.5498 | 0.391248 |
| **HTR2C** | 0.00113103 | 0.00488147 | **Mrgprg** | NA | NA |
| **HTR4** | 0.00271656 | 0.00759749 | **Mrgprx1** | NA | 0.0105185 |
| **HTR5A** | NA | NA | **Mrgprx2** | 0.0774361 | 12.2995 |
| **HTR6** | 0.087829 | 0.0632873 | **Mtnr1a** | NA | 0.00719732 |
| **HTR7** | 0.0644815 | 0.0179197 | **Mtnr1b** | NA | 0.0236863 |
| **KISS1R** | 0.0991392 | NA | **Nmbr** | 1.50229 | 49.4062 |
| **LGR4** | 3.37246 | 15.8887 | **Nmur1** | 0.0185323 | 0.0209506 |
| **LGR5** | 0.209288 | 0.00130332 | **Nmur2** | NA | NA |
| **LGR6** | 0.00878684 | 0.273049 | **Npbwr1** | NA | 0.0188803 |
| **LHCGR** | 0.0115847 | 0.00276917 | **Npffr1** | 0.0111858 | 0.0358288 |
| **LPAR1** | 0.30614 | 0.00154123 | **Npffr2** | NA | 0.0123785 |
| **LPAR2** | 0.163059 | 2.24396 | **Npsr1** | 0.00677573 | 0.00414911 |
| **LPAR3** | 1.02485 | NA | **Npy1r** | 0.924803 | 0.0435617 |
| **LPAR4** | NA | NA | **Npy2r** | NA | 0.136509 |
| **LPAR5** | 0.0071762 | 2.32662 | **Npy4r** | 0.203163 | NA |
| **LPAR6** | 0.454705 | 0.349783 | **Npy5r** | 0.0302107 | 0.0341529 |
| **LTB4R** | NA | NA | **Npy6r** | NA | NA |
| **LTB4R** | 4.66447 | 13.2965 | **Ntsr1** | 0.00274014 | 0.00154885 |
| **LTB4R2** | NA | NA | **Ntsr2** | 0.108081 | 0.336006 |
| **LTB4R2** | 19.3295 | 23.0131 | **Opn1mw** | 0.0211735 | 8.35383 |
| **MAS1** | NA | 0.00721669 | **Opn1sw** | 0.742135 | 0.0886244 |
| **MAS1L** | NA | NA | **Opn3** | 3.99339 | 1.40762 |
| **MAS1L** | NA | NA | **Opn4** | 0.103576 | NA |
| **MAS1L** | NA | NA | **Opn5** | NA | 0.00765818 |
| **MAS1L** | NA | NA | **Oprd1** | 0.0255691 | 0.251892 |
| **MAS1L** | NA | NA | **Oprk1** | NA | 0.0315662 |
| **MAS1L** | NA | NA | **Oprl1** | 0.0413132 | 2.05498 |
| **MAS1L** | NA | NA | **Oprm1** | 0.00264547 | 0.016698 |
| **MAS1L** | NA | NA | **Oxgr1** | NA | NA |
| **MC1R** | 1.04787 | 10.2072 | **Oxtr** | 0.0455027 | 0.0600138 |
| **MC2R** | NA | NA | **P2ry1** | NA | 0.0225294 |
| **MC3R** | NA | NA | **P2ry10** | 0.0149633 | 0.0422896 |
| **MC4R** | 0.143407 | NA | **P2ry12** | 0.00264635 | 0.0254292 |
| **MC5R** | NA | NA | **P2ry13** | NA | 0.163137 |
| **MCHR1** | 0.0302336 | NA | **P2ry14** | 0.0248104 | 0.00175299 |
| **MCHR2** | 0.0056801 | 0.000840513 | **P2ry2** | 3.9949 | 0.359243 |
| **MLNR** | 0.150612 | 0.124806 | **P2ry4** | 0.0522985 | NA |

| | | | | | |
|---|---|---|---|---|---|
| **MRGPRD** | NA | 0.263566 | **P2ry6** | 0.103266 | 0.176995 |
| **MRGPRE** | NA | 0.0344619 | **Prlhr** | NA | 0.0443141 |
| **MRGPRF** | NA | 0.899388 | **Prokr1** | 0.629131 | 0.69402 |
| **MRGPRG** | NA | NA | **Prokr2** | NA | 0.0701224 |
| **MRGPRX1** | NA | NA | **Ptafr** | 0.119233 | 0.0299537 |
| **MRGPRX2** | NA | NA | **Ptgdr** | NA | NA |
| **MRGPRX3** | 0.0105186 | 0.00726363 | **Ptgdr2** | 1.08297 | 0.481899 |
| **MRGPRX4** | NA | NA | **Ptger1** | 39.192 | 10.0268 |
| **MTNR1A** | 0.0813236 | NA | **Ptger2** | 0.181515 | NA |
| **MTNR1B** | 0.0119583 | 0.0206445 | **Ptger3** | 0.24821 | 0.0070591 |
| **NMBR** | 0.598969 | 0.515978 | **Ptger4** | 8.76749 | 2.31443 |
| **NMUR1** | 0.118515 | NA | **Ptgfr** | 1.55615 | 0.0152533 |
| **NMUR2** | 0.0293856 | 0.00608767 | **Ptgir** | 16.0573 | 39.3517 |
| **NPBWR1** | 0.0944758 | 0.12582 | **Pth1r** | 0.167307 | 0.111258 |
| **NPBWR2** | NA | NA | **Pth2r** | 0.0207088 | 0.0565769 |
| **NPBWR2** | NA | NA | **Qrfpr** | 0.0172516 | 0.0162523 |
| **NPFFR1** | NA | 0.024297 | **Rho** | 0.743877 | 0.176599 |
| **NPFFR2** | 0.0269554 | 0.0136868 | **Rxfp1** | 0.00256196 | 0.0101369 |
| **NPSR1** | 0.00139586 | 0.000289172 | **Rxfp2** | NA | 0.00878048 |
| **NPY1R** | 0.176656 | NA | **Rxfp3** | 0.144234 | 0.0163055 |
| **NPY2R** | 0.116451 | NA | **Rxfp4** | 0.891469 | 0.111978 |
| **NPY4R** | 0.037932 | 14.8781 | **S1pr1** | 46.7234 | 0.600914 |
| **NPY5R** | 0.0382008 | NA | **S1pr2** | 12.391 | 2.64884 |
| **NPY6R** | 0.179997 | 0.173128 | **S1pr3** | 11.6448 | 0.0443078 |
| **NTSR1** | 0.00683586 | 2.96093 | **S1pr4** | NA | 0.116858 |
| **NTSR2** | 0.00511315 | 0.0211853 | **S1pr5** | 0.156045 | 0.113405 |
| **OPN1LW** | 0.0207741 | NA | **Sctr** | 0.0892628 | 0.980804 |
| **OPN1MW** | 0.0215371 | NA | **Smo** | 14.9102 | 0.0889417 |
| **OPN1SW** | 10.4651 | 18.0731 | **Sstr1** | NA | 0.0260778 |
| **OPN3** | 12.3259 | 13.9307 | **Sstr2** | NA | 1.02156 |
| **OPN4** | NA | 0.0103481 | **Sstr3** | 0.0803073 | 0.508405 |
| **OPN5** | NA | NA | **Sstr4** | NA | 0.195803 |
| **OPRD1** | 0.226568 | 0.0150715 | **Sstr5** | NA | NA |
| **OPRK1** | NA | NA | **Sucnr1** | NA | NA |
| **OPRL1** | NA | NA | **Taar1** | NA | NA |
| **OPRL1** | 0.116552 | 0.238359 | **Taar2** | NA | NA |
| **OPRM1** | 0.00935894 | 0.00403926 | **Taar5** | NA | NA |
| **OR1A1** | NA | NA | **Taar6** | NA | NA |
| **OR1G1** | NA | NA | **Taar9** | NA | NA |
| **OR2T11** | NA | NA | **Tacr1** | NA | 0.00441818 |
| **OR51E1** | NA | NA | **Tacr2** | 0.0272973 | 0.072005 |
| **OXER1** | 0.207018 | NA | **Tacr3** | 0.0046724 | 0.02509 |
| **OXGR1** | 0.0477306 | NA | **Tas1r1** | 0.694434 | 0.588788 |
| **OXTR** | 0.479552 | 1.0332 | **Tas1r2** | 0.0515582 | NA |
| **P2RY1** | 19.3826 | NA | **Tas1r3** | 1.95746 | 0.476622 |
| **P2RY10** | NA | NA | **Tbxa2r** | 0.268003 | 0.205591 |
| **P2RY11** | 27.7526 | 32.5348 | **Tpra1** | 15.236 | 4.63404 |
| **P2RY12** | 0.98374 | NA | **Trhr** | 0.0248016 | 0.0122666 |
| **P2RY13** | 0.189777 | NA | **Tshr** | 0.0246273 | 0.0236226 |
| **P2RY14** | 0.442191 | 0.00287468 | **Uts2r** | 0.0724133 | 0.0409313 |
| **P2RY2** | 0.0865239 | 2.76515 | **Vipr1** | 0.024469 | 0.0345775 |
| **P2RY4** | 0.11558 | 0.0399067 | **Vipr2** | 0.16374 | 0.0122049 |

| | | | | | |
|---|---|---|---|---|---|
| **P2RY6** | NA | 2.84855 | **Xcr1** | 0.0126938 | NA |
| **P2RY8** | 0.00164734 | 0.0307143 | | | |
| **PRLHR** | NA | NA | | | |
| **PROKR1** | 0.0376915 | NA | | | |
| **PROKR2** | 0.0249188 | NA | | | |
| **PTAFR** | 0.110363 | 0.187806 | | | |
| **PTGDR** | NA | NA | | | |
| **PTGDR2** | 3.83296 | 5.86027 | | | |
| **PTGER1** | 0.0212703 | 0.308452 | | | |
| **PTGER2** | 0.133269 | 0.035624 | | | |
| **PTGER3** | 0.0154052 | 0.00293089 | | | |
| **PTGER4** | 0.182291 | 4.79882 | | | |
| **PTGFR** | 1.00315 | 0.0142452 | | | |
| **PTGIR** | NA | 0.0275427 | | | |
| **PTH1R** | 0.0683325 | 0.0439327 | | | |
| **PTH2R** | 0.00248387 | 0.00192964 | | | |
| **QRFPR** | 0.219221 | NA | | | |
| **RHO** | 0.0091634 | NA | | | |
| **RXFP1** | 0.00272656 | 0.0826561 | | | |
| **RXFP2** | 0.0019244 | NA | | | |
| **RXFP3** | NA | NA | | | |
| **RXFP3** | NA | NA | | | |
| **RXFP4** | 0.296654 | 0.965742 | | | |
| **S1PR1** | 1.12151 | 2.05386 | | | |
| **S1PR2** | 1.48363 | 1.69754 | | | |
| **S1PR3** | 1.08342 | 2.00844 | | | |
| **S1PR4** | 0.0153875 | 1.41057 | | | |
| **S1PR5** | 0.296001 | NA | | | |
| **SCTR** | 0.0287622 | 0.00372408 | | | |
| **SMO** | 6.11217 | NA | | | |
| **SSTR1** | 0.0356989 | 9.25679 | | | |
| **SSTR2** | 1.50664 | 0.109517 | | | |
| **SSTR3** | NA | 0.0154474 | | | |
| **SSTR4** | NA | NA | | | |
| **SSTR5** | NA | 3.13115 | | | |
| **SUCNR1** | NA | NA | | | |
| **TAAR1** | NA | NA | | | |
| **TAAR2** | NA | NA | | | |
| **TAAR3P** | NA | NA | | | |
| **TAAR5** | NA | NA | | | |
| **TAAR6** | NA | NA | | | |
| **TAAR8** | NA | NA | | | |
| **TAAR9** | NA | NA | | | |
| **TACR1** | 0.169425 | 0.218328 | | | |
| **TACR2** | 0.0236055 | 0.948704 | | | |
| **TACR3** | 0.00275241 | 0.00997854 | | | |
| **TAS1R1** | 0.0502677 | 0.0442583 | | | |
| **TAS1R2** | 0.00611928 | 0.025354 | | | |
| **TAS1R3** | 0.609168 | 2.96566 | | | |
| **TAS2R1** | 0.00147393 | 0.0183208 | | | |
| **TAS2R10** | NA | NA | | | |
| **TAS2R10** | NA | NA | | | |

| | | |
|---|---|---|
| **TAS2R10** | 1.2974 | 1.83257 |
| **TAS2R13** | NA | NA |
| **TAS2R13** | NA | NA |
| **TAS2R13** | 1.31383 | 0.466594 |
| **TAS2R14** | NA | NA |
| **TAS2R14** | NA | NA |
| **TAS2R14** | 1.43412 | 0.57028 |
| **TAS2R16** | NA | NA |
| **TAS2R19** | NA | NA |
| **TAS2R19** | NA | NA |
| **TAS2R19** | 3.6183 | 3.9385 |
| **TAS2R20** | NA | NA |
| **TAS2R20** | NA | NA |
| **TAS2R20** | 3.60422 | 3.64116 |
| **TAS2R3** | 1.56276 | 0.404685 |
| **TAS2R30** | NA | NA |
| **TAS2R30** | NA | NA |
| **TAS2R30** | 2.76834 | 0.113191 |
| **TAS2R31** | NA | NA |
| **TAS2R31** | NA | NA |
| **TAS2R31** | 2.40744 | 1.1845 |
| **TAS2R38** | 0.0537618 | NA |
| **TAS2R39** | 0.120845 | NA |
| **TAS2R4** | 2.69409 | 0.202953 |
| **TAS2R40** | 0.294582 | NA |
| **TAS2R41** | NA | NA |
| **TAS2R41** | NA | NA |
| **TAS2R42** | NA | NA |
| **TAS2R42** | 0.260105 | NA |
| **TAS2R43** | NA | NA |
| **TAS2R43** | NA | NA |
| **TAS2R43** | NA | NA |
| **TAS2R45** | NA | NA |
| **TAS2R46** | NA | NA |
| **TAS2R46** | NA | NA |
| **TAS2R46** | 2.24655 | 0.342211 |
| **TAS2R5** | 2.93635 | 0.497707 |
| **TAS2R50** | NA | NA |
| **TAS2R50** | NA | NA |
| **TAS2R50** | 1.84349 | 0.827465 |
| **TAS2R60** | NA | NA |
| **TAS2R60** | NA | NA |
| **TAS2R7** | NA | NA |
| **TAS2R7** | NA | NA |
| **TAS2R7** | NA | NA |
| **TAS2R8** | NA | NA |
| **TAS2R8** | NA | NA |
| **TAS2R8** | 0.197747 | NA |
| **TAS2R9** | NA | NA |
| **TAS2R9** | NA | NA |
| **TAS2R9** | 0.0571626 | NA |
| **TBXA2R** | 0.0397444 | 3.14422 |

| | | |
|---|---|---|
| **TPRA1** | 3.62526 | 7.53851 |
| **TRHR** | 0.11594 | 0.0564074 |
| **TSHR** | 0.0234475 | 0.00532329 |
| **UTS2R** | NA | NA |
| **VIPR1** | 0.392057 | 0.520812 |
| **VIPR2** | 0.0189574 | 0.147819 |
| **XCR1** | 0.028664 | NA |

**Table S5. Primer Sequences**

| Primer Name | Corresponding cDNA Name | Sequence |
| --- | --- | --- |
| CDH-ADRB2F | β2-AR | GTCGTGATCTAGAGCTAGCG CACC ATGGGGCAACCCGGGAACGGCAGC |
| CDH-ADRB2R | β2-AR | CCAGAGGTTGATTGTCGACGC TTACAGCAGTGAGTCATTTGTACTACAATTCCTCCCTTG |
| CIG-GNAQF | Gaq | TCGAGCTCAAGCTTCGCACC ATGACTCTGGAGTCCATCATGGCG |
| CIG-GNAQR | Gaq | TAGAAGCTTCTGCAGAT TTAGACCAGATTGTACTCCTTCAGGTTCAAC |
| Q_Q209LR | Gaq CA | CAGGCCCCCTACATCGACCATTCTG |
| Q_Q209LF | Gaq CA | TGGTCGATGTAGGGGGCCTGAGGTCAGAGAGAAGAAAATGGATACACTGC |
| CIG-GNASF | Gas | TCGAGCTCAAGCTTCGCACC ATGGGCTGCCTCGGGAACAGTAAG |
| CIG-GNASR | Gas | TAGAAGCTTCTGCAGAT TTAGAGCAGCTCGTACTGACGAAGGTG |
| S_Q227LR | Gas CA | TCATCGCGCAGGCCACCCACGTCAAACATGT |
| S_Q227LF | Gas CA | GACGTGGGTGGCCTGCGCGATGAACGCCGCAAG |
| CIG-GNAi2F | Gai | TCGAGCTCAAGCTTCGCACC ATGGGCTGCACCGTGAGCG |
| CIG-GNAi2R | Gai | TAGAAGCTTCTGCAGAT TCAGAAGAGGCCGCAGTCCTTCAGGTTG |
| i2_Q205LF | Gai CA | GTGGGTGGTCTGCGGTCTGAGCGG |
| i2_Q205LR | Gai CA | CCGCTCAGACCGCAGACCACCCAC |
| CIG-EPAC2F | EPAC | TCGAGCTCAAGCTTCGCACC ATGGTCGCTGCGCACGCTG |
| CIG-EPAC2R | EPAC | TAGAAGCTTCTGCAGAT CTATGGTCGACGAGGCTCTAATCTGTG |
| R432KF | EPAC DN | CTAGTGAATGATGCCCCAAAGGCTGCCTCTATCGTCTTA |
| R432KR | EPAC DN | TAAGACGATAGAGGCAGCCTTTGGGGCATCATTCACTAG |
| CIG-RAP1AS17NF | RAP1A DN | TCGAGCTCAAGCTTCGCACC ATGCGTGAGTACAAGCTAGTGGTCCTTGGTTCAGGAGGCGTTGGGAAGAATGCTCTGACAGTTCAGTTTG |
| CIG-RAP1AS17NR | RAP1A DN | TAGAAGCTTCTGCAGAT CTAGAGCAGCAGACATGATTTCTTTTTAGGCTTCTTCTTTTCC |
